# SARS-CoV-2 papain-like protease PLpro in complex with natural compounds reveal allosteric sites for antiviral drug design

**DOI:** 10.1101/2021.11.17.468943

**Authors:** Vasundara Srinivasan, Hévila Brognaro, Prince R. Prabhu, Edmarcia Elisa de Souza, Sebastian Günther, Patrick Y. A. Reinke, Thomas J. Lane, Helen Ginn, Huijong Han, Wiebke Ewert, Janina Sprenger, Faisal H. M. Koua, Sven Falke, Nadine Werner, Hina Andaleeb, Najeeb Ullah, Bruno Alves Franca, Mengying Wang, Angélica Luana C Barra, Markus Perbandt, Martin Schwinzer, Christina Schmidt, Lea Brings, Kristina Lorenzen, Robin Schubert, Rafael Rahal Guaragna Machado, Erika Donizette Candido, Danielle Bruna Leal Oliveira, Edison Luiz Durigon, Oleksandr Yefanov, Julia Lieske, Luca Gelisio, Martin Domaracky, Philipp Middendorf, Michael Groessler, Fabian Trost, Marina Galchenkova, Sofiane Saouane, Johanna Hakanpää, Markus Wolf, Dusan Turk, Arwen R. Pearson, Henry N. Chapman, Winfried Hinrichs, Carsten Wrenger, Alke Meents, Christian Betzel

## Abstract

SARS-CoV-2 papain-like protease (PLpro) covers multiple functions. Beside the cysteine-protease activity, PLpro has the additional and vital function of removing ubiquitin and ISG15 (Interferon-stimulated gene 15) from host-cell proteins to aid coronaviruses in evading the host’s innate immune responses. We established a high-throughput X-ray screening to identify inhibitors by elucidating the native PLpro structure refined to 1.42 Å and performing co-crystallization utilizing a diverse library of selected natural compounds. We identified three phenolic compounds as potential inhibitors. Crystal structures of PLpro inhibitor complexes, obtained to resolutions between 1.7-1.9 Å, show that all three compounds bind at the ISG15/Ub-S2 allosteric binding site, preventing the essential ISG15-PLpro molecular interactions. All compounds demonstrate clear inhibition in a deISGylation assay, two exhibit distinct antiviral activity and one inhibited a cytopathic effect in a non-cytotoxic concentration range. These results highlight the druggability of the rarely explored ISG15/Ub-S2 PLpro allosteric binding site to identify new and effective antiviral compounds. Importantly, in the context of increasing PLpro mutations in the evolving new variants of SARS-CoV-2, the natural compounds we identified may also reinstate the antiviral immune response processes of the host that are down-regulated in COVID-19 infections.

## Introduction

After more than one and half year, the coronavirus disease COVID-19, caused by SARS-CoV-2, remains devastating with high numbers of infections and deaths^1^. Several approved and commercialized vaccines against COVID-19 were developed worldwide in only a year. However, these vaccines are not uniformly available around the world, and consequently new SARS-CoV-2 variants have already emerged which may impact the effectiveness of the available vaccines in the future^2^. In parallel, more efforts are needed to identify and optimize alternative treatments for patients infected by SARS-CoV-2, who do not respond to or cannot tolerate vaccines^3^. Hence, research is ongoing at a rapid pace to identify effective drug candidates by applying complementary strategies. One approach, we followed recently, is the massive X-ray crystallographic screening for inhibitors of SARS-CoV-2 main protease Mpro, an essential protein in the viral replication process and hence an important drug target^4^. We identified six compounds inhibiting Mpro that showed antiviral activity and these compounds are currently approaching the step of pre-clinical investigations^4^.

The positive-sense single stranded RNA genome of coronaviruses encodes 16 nonstructural polyproteins (nsps 1-16). The Papain-like protease (PLpro) is a domain that is part of the nsp3 gene, the largest mature SARS-CoV-2 protein^5^. PLpro is required to recognize and cleave the motif LXGG within preprocessed polyproteins between nsp1/2, nsp2/3 and nsp3/4 into functional units for initiation, replication and transcription of the viral genome^6, 7^. Apart from the proteolytic activity, PLpro can also bind and cleave ubiquitin chains or ISG15 (interferon-stimulated gene product 15) from ubiquitinated or ISGylated proteins^8^. The deubiquitinase (DUB), as well as deISGylating activities are vital for the coronaviruses to antagonize the host immune responses. It has been shown that mono-ubiquitination at the endoplasmic reticulum (ER) membranes regulate endocytosis, and vesicle trafficking and thus is important for coronavirus propagation^9^. On the other hand, ISG15 causes metabolic pathway modifications towards excessive inflammatory and autoimmune responses due to interferon system dysregulations^10, 11^. Target proteins for the coronavirus, such as IRF3 (Interferon regulatory factor 3) need to conjugate with ISG15 to be correctly phosphorylated for their entry into nucleus where they are required for downstream signalling events for e.g., *via* the IFN-β pathway^12^ to elicit an antiviral immune response. Therefore, the cysteine protease activity, together with the deubiquitinase and deISGylating activity of PLpro undoubtedly makes this enzyme a very promising target for drug discovery investigations^20, 21^. Furthermore, the importance of PLpro as a drug target was highlighted in several recent studies that have identified novel and unexpected mutations in the PLpro domain of the nsp3 gene from the SARS-CoV-2 variants of concern (VOC), currently circulating in different parts of the world^13, 14^.

The crystal structure of SARS-CoV PLpro, in complex with Lys48-linked di-ubiquitin^15^ (PDB code 5E6J), clearly evokes a mechanism of DUB action in which the LXGG residues at the C-terminal of the ubiquitin molecule bind in a cleft (Ub S1 proximal binding site) located close to the catalytic active site, allowing efficient cleavage. In addition, the recently obtained crystal structure of SARS-CoV-2 PLpro, in complex with mouse-ISG15^16^, demonstrates that the N-terminal of ISG15 interacts with PLpro at a different binding site termed as ISG15/Ub S2 distal binding site. Compared to the crystal structure of ISG15 (PDB code 5TLA), the N-terminal half of the ISG15 molecule is rotated by about 90° and coordinates with the S2-helix, as shown in the complex structure of SARS-CoV-2 PLpro with ISG15.

These different binding events explain and highlight why SARS-CoV-2 PLpro and SARS-CoV PLpro, despite sharing 83% sequence identity show different host substrate preferences. Recently data were published showing higher affinity and specificity of SARS-CoV-2 PLpro to ISG15, whereas SARS-CoV PLpro preferentially cleaves ubiquitin chains, which may be associated to the substantial higher morbidity and mortality of SARS-CoV-2 in comparison to SARS-CoV infections^17^.

Until now several high throughput assay screening (HTS) activities, focussed on identifying potential inhibitors of PLpro using repurposing compound libraries were initiated. Unfortunately, these were not very successful in obtaining hits of compounds that could be further developed to obtain an effective antiviral drug^17^. Hence, we set out a different strategy to explore a unique library consisting of 500 natural compounds assembled and characterized by the Molecular Bank, ICCBS, Karachi, Pakistan, by structure-based drug design (https://iccs.edu/page-mol-bank). Natural compounds of the ICCBS Molecular Bank extracted from plants present characteristics such as high chemical diversity, medically relevant anti-tumor, anti-oxidant, anti-inflammatory and importantly antiviral action with typically milder or no side effects, in addition to lower cost of production as compared to most available drugs on the market^18^. A number of these compounds also have a long history of use as drug molecules to treat distinct human diseases, including viral infections, such as Hepatitis C virus infection^19^. Recent reports also demonstrate the potential use of plant molecules and their secondary metabolites against SARS-CoV-2, and other human coronaviruses^20–22^ by applying *in vitro* and *in silico* approaches. However, to our knowledge, a systematic screening of natural products by structure-based drug discovery providing direct experimental data about complex formation was not available to date.

Our high throughput screening by X-ray crystallography identified three natural compounds bound to PLpro. All three compounds, 4-(2- hydroxyethyl)phenol (YRL), 4-hydroxybenzaldehyde (HBA) and methyl 3, 4- dihydroxybenzoate (HE9) are polyphenols, one of the most important and largest class of bioactive compounds present in plants. This vast group of bioactive compounds is divided into six major classes: hydroxybenzoic acids, hydroxycinnamic acids, flavonoids, stilbenes, and lignans. In addition to the well-known anti-oxidant and anti-inflammatory activities of polyphenols, several studies have reported their anti-viral potential against Epstein-Barr virus^23, 24^, enterovirus 71^25^, herpes simplex virus (HSV)^26^, influenza virus^27^, and other viruses causing respiratory tract-related infections^28^.

Interestingly, all three compounds YRL, HBA and HE9 bind at the same, and yet unexplored ISG15/Ub-S2 allosteric binding site of PLpro by forming specific interactions and clearly inhibit PLpro in deISGylation activity assays. None of these lead compounds are cytotoxic in cellular cytotoxicity assays, demonstrating their potential as drug molecules. Significantly, two of them exhibit antiviral activity and one inhibits cytopathic effects in the range of 60-80% in a non-cytotoxic concentration range up to 100 µM in cellular assays. These three natural phenolic compounds undoubtedly provide a scaffold as antiviral drugs for further development and optimization towards the prevention and/or reduction of SARS-CoV-2 viral replication, and to reinstate and support the innate immune response of the host in parallel.

## Results

### X-ray screening of a natural compound library identifies three allosteric inhibitors of SARS-CoV-2 PLpro

We initiated a structure-based drug discovery approach to identify potential inhibitors for PLpro by X-ray screening of 500 compounds from the ICCBS Molecular Bank. SARS-CoV-2 PLpro was expressed recombinantly in *Escherichia coli*, and purified to homogeneity as a monomer (see materials and methods). Wild-type enzyme crystals were obtained in a stable and reproducible condition and diffracted X-rays to a high resolution of 1.42 Å. Data collection and refinement statistics are summarized in the Supplementary Table 1. The electron density maps obtained for the wild-type enzyme allowed the elucidation of all 315 amino acid residues, the zinc ion and 529 solvent water molecules. Further, a glycerol molecule from the cryoprotectant, used for freezing crystals, could be modelled in the electron density map, as well as a phosphate and two chloride ions.

PLpro folds with a right-handed architecture consisting of thumb, palm, and fingers domains with a catalytic triad consisting of Cys111-His272-Asp286 and a N-terminal ubiquitin-like domain (Figure 1). Four cysteine side chains coordinate a zinc ion, constituting a ‘zinc finger motif’ that is essential for structural stability and protease activity of the enzyme^29^. The overall structure of SARS-CoV-2 PLpro is similar to SARS-CoV PLpro (PDB code 2FE8) that shares a sequence identity of 83% with a r.m.s.d. of 0.58Å for 260 equivalent Cα atoms and also to MERS-CoV PLpro (PDB code 4RNA) despite a lower sequence identity of 29% and a corresponding r.m.s.d. of 1.83Å for 258 equivalent Cα atoms (Figure S1 & S3). The most structurally dynamic regions are the ubiquitin-fold like, and the zinc fingers domains. The catalytic active site region is conformationally well conserved among the different coronaviral PLpro enzymes. The access to the active site is regulated *via* a flexible loop named “blocking loop 2” (BL2, Figure 1), as this loop changes from an ‘open’ to a ‘closed’ conformation in the context of substrate binding^30^. A number of known PLpro inhibitors bind at this site, including the high affinity inhibitor GRL0617, and structural variations have been observed in this loop among different PLpro enzymes^31^.

**Figure 1:**
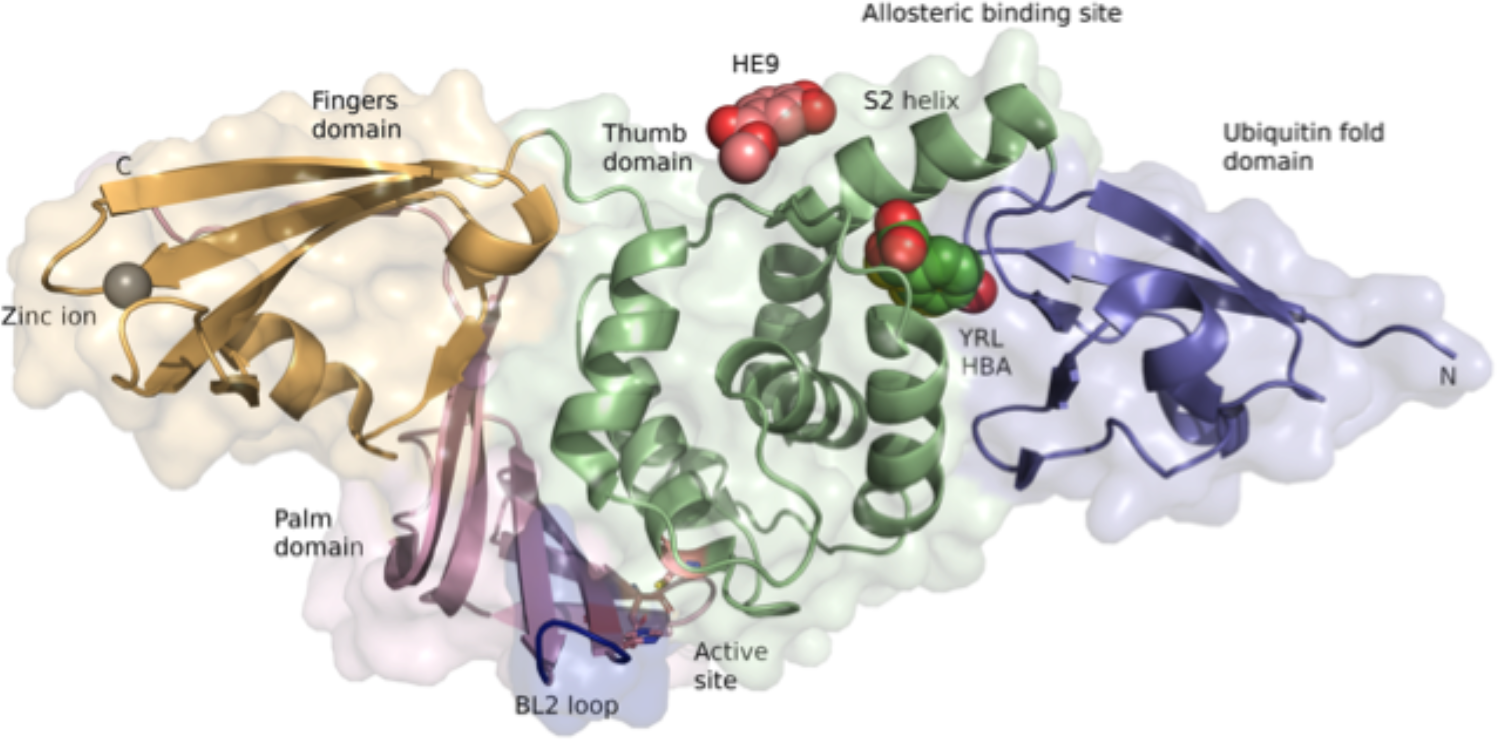
Crystal structures of SARS-CoV-2 PLpro complexes with the three natural compounds. PLpro domains are depicted in a right-handed architecture, ubiquitin-fold like (blue), thumb (green), palm (salmon pink) and fingers (light orange). Catalytic active site residues Cys 111, His 272 and Asp 286 are represented as sticks and a zinc ion in the fingers domain is shown as a grey sphere. The flexible blocking loop (BL2 loop) that changes conformation in the context of substrate binding is shown in blue. YRL (green spheres), HBA (yellow spheres) and HE9 (pink spheres) compounds bind at the allosteric site that is located about 30 Å apart to the active site. S2 helix involved in the interaction of the ISG15 molecule is indicated.

Crystals of SARS-CoV-2 PLpro, in complex with the three natural compounds, were obtained by co-crystallization using the vapour diffusion method in a screening approach utilizing 500 molecules from a library of natural compounds. Crystals were grown in the same condition as for the native PLpro and diffraction datasets were collected in the resolution range of 1.7-1.9 Å. Over 2,000 crystals were harvested, and multiple datasets for each compound were collected that resulted in ∼2500 diffraction datasets. PLpro structures complexed with inhibitor compounds were solved using the ligand-free PLpro (PDB code 7NFV) as reference model for consistent indexing of datasets with a previously established automatic pipeline^4^ (see materials and methods). Data collection and refinement statistics are summarized in Table 1. The three complex structures obtained superimpose well with the ligand-free structure 7NFV with a r.m.s.d. of 0.26 Å (298 Cα atoms) to 7OFS, r.m.s.d. of 0.07 Å (283 Cα atoms) to 7OFU and r.m.s.d. of 0.33 Å (299 Cα atoms) to 7OFT, respectively. The three compounds bind at the same ISG15/Ub S2 allosteric site in PLpro (Figure 1) that is located about 30 Å apart to the active site residue Cys 111. The interaction between these allosteric inhibitors and PLpro are formed *via* hydrogen bonds, hydrophobic, and π-stacking interactions (Figure S2A, B, C).

**Table 1.**
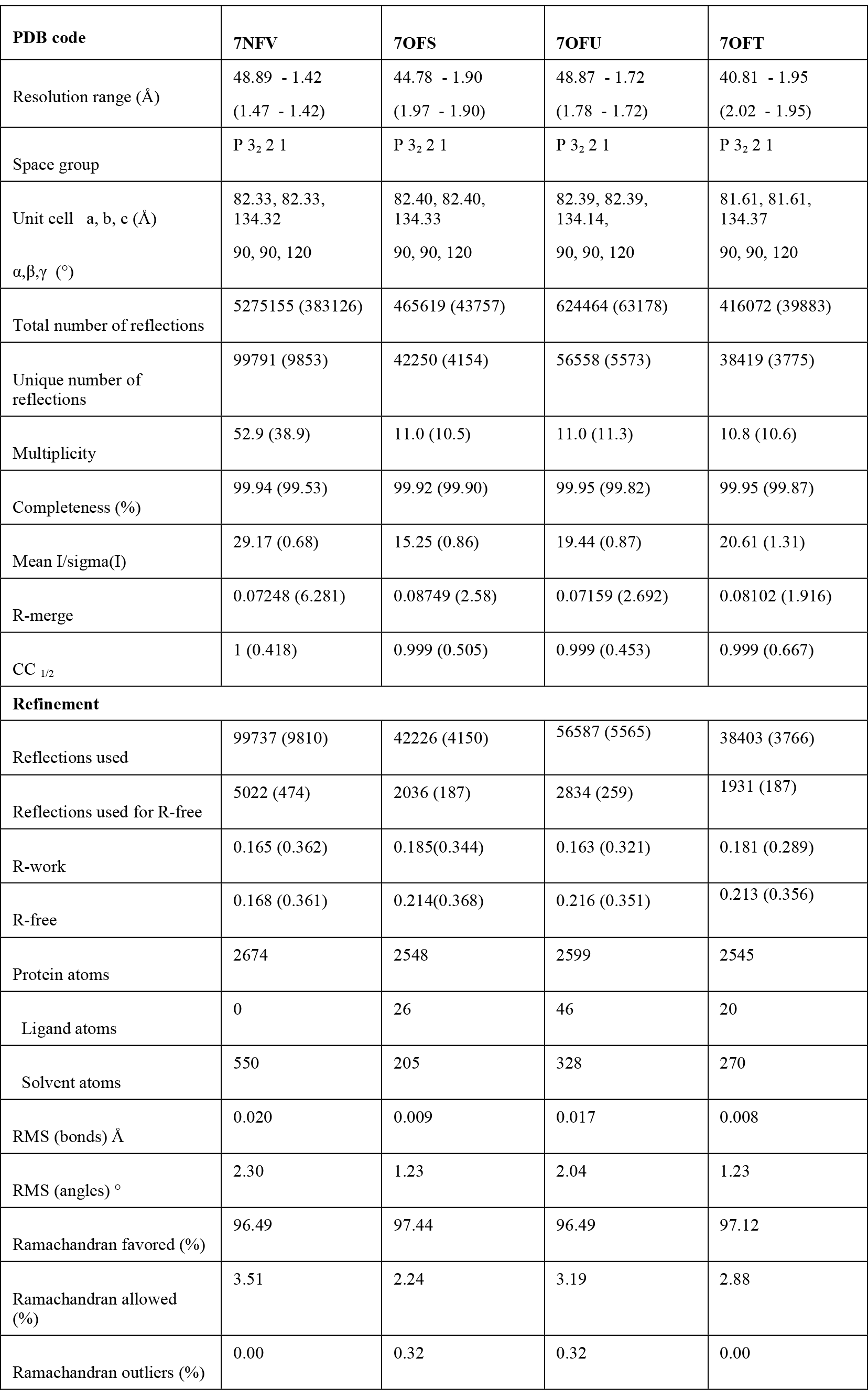
Data collection and refinement statistics

| PDB code | 7NFV | 7OFS | 7OFU | 7OFT |
| --- | --- | --- | --- | --- |
| Resolution range (Å) | 48.89 - 1.42<br>(1.47 - 1.42) | 44.78 - 1.90<br>(1.97 - 1.90) | 48.87 - 1.72<br>(1.78 - 1.72) | 40.81 - 1.95<br>(2.02 - 1.95) |
| Space group | P 3<sub>2</sub> 2 1 | P 3<sub>2</sub> 2 1 | P 3<sub>2</sub> 2 1 | P 3<sub>2</sub> 2 1 |
| Unit cell a, b, c (Å) | 82.33, 82.33,<br>134.32 | 82.40, 82.40,<br>134.33 | 82.39, 82.39,<br>134.14, | 81.61, 81.61,<br>134.37 |
| $\alpha, \beta, \gamma$ (°) | 90, 90, 120 | 90, 90, 120 | 90, 90, 120 | 90, 90, 120 |
| Total number of reflections | 5275155 (383126) | 465619 (43757) | 624464 (63178) | 416072 (39883) |
| Unique number of reflections | 99791 (9853) | 42250 (4154) | 56558 (5573) | 38419 (3775) |
| Multiplicity | 52.9 (38.9) | 11.0 (10.5) | 11.0 (11.3) | 10.8 (10.6) |
| Completeness (%) | 99.94 (99.53) | 99.92 (99.90) | 99.95 (99.82) | 99.95 (99.87) |
| Mean I/sigma(I) | 29.17 (0.68) | 15.25 (0.86) | 19.44 (0.87) | 20.61 (1.31) |
| R-merge | 0.07248 (6.281) | 0.08749 (2.58) | 0.07159 (2.692) | 0.08102 (1.916) |
| CC<sub>1/2</sub> | 1 (0.418) | 0.999 (0.505) | 0.999 (0.453) | 0.999 (0.667) |
| **Refinement** |  |  |  |  |
| Reflections used | 99737 (9810) | 42226 (4150) | 56587 (5565) | 38403 (3766) |
| Reflections used for R-free | 5022 (474) | 2036 (187) | 2834 (259) | 1931 (187) |
| R-work | 0.165 (0.362) | 0.185(0.344) | 0.163 (0.321) | 0.181 (0.289) |
| R-free | 0.168 (0.361) | 0.214(0.368) | 0.216 (0.351) | 0.213 (0.356) |
| Protein atoms | 2674 | 2548 | 2599 | 2545 |
| Ligand atoms | 0 | 26 | 46 | 20 |
| Solvent atoms | 550 | 205 | 328 | 270 |
| RMS (bonds) Å | 0.020 | 0.009 | 0.017 | 0.008 |
| RMS (angles) ° | 2.30 | 1.23 | 2.04 | 1.23 |
| Ramachandran favored (%) | 96.49 | 97.44 | 96.49 | 97.12 |
| Ramachandran allowed (%) | 3.51 | 2.24 | 3.19 | 2.88 |
| Ramachandran outliers (%) | 0.00 | 0.32 | 0.32 | 0.00 |

### Molecular basis of inhibition by the three allosteric inhibitors of SARS-CoV-2 PLpro

The natural compound 4-(2-hydroxyethyl)phenol (YRL), isolated from *Lawsonia alba*, is a well-known antioxidant and an anti-arrythmia agent (https://pubchem.ncbi.nlm.nih.gov/compound/2-_4-Hydroxyphenyl_ethanol). YRL binds to PLpro at the ISG15/Ub S2 allosteric binding site in a hydrophobic cavity with a predicted binding energy of -7.17 kcal/mol (calculated using Prodigy^32^). The benzene core is covered by hydrophobic interactions with side chains of Val 11, Val 57, Pro 59, Tyr 72 and Leu 80. The main-chain nitrogen atom of Leu 80 is hydrogen bonded *via* a water molecule to the hydroxyethyl substituent of YRL. Interestingly, the hydroxyethyl substituent is observed with two alternative conformations and is refined to equal occupancy in the complex structure. One conformation forms a hydrogen bond to carbonyl backbone of Asp 76, the alternative hydroxyl to the carbonyl of Thr 74. Both alternative hydroxyl groups replace water molecules in the ligand-free enzyme and have contacts to solvent water molecules at the entry of the binding pocket. The phenolic hydroxyl is hydrogen bonded to carbonyl oxygen of Val 57 at the bottom of the binding pocket to complete the interaction of the ligand YRL in the PLpro-YRL complex structure (Figure S2A).

The second compound, 4-hydroxybenzaldehyde (HBA), isolated from *Acalypha torta*, is a well-known anti-tumor agent devoid of any side effects (https://pubchem.ncbi.nlm.nih.gov/compound/4-Hydroxybenzaldehyde). The calculated binding energy for the interaction of the HBA ligand to PLpro is -6.97 kcal/mol. The benzene core and the phenolic hydroxyl is observed in the same position and, thus, has similar interaction to PLpro as described above for YRL (Figure S2B). The distance of the phenolic hydroxyl of both compounds to the Cα of Val 11 indicates a C-H···O hydrogen bond (3.4 Å). The aldehyde substituent has weak water contacts at the entrance of the binding pocket. The remarkable structural change in PLpro to accommodate these two compounds is that the side chain of Leu 80 has to tilt away in the complex PLpro structures (Figure S8).

The third compound, methyl 3,4-dihydroxybenzoate (HE9), isolated from *Tagetes patula* (marigold), is a major diphenol found in green tea with antioxidant and anti-inflammatory effects^33^ (https://pubchem.ncbi.nlm.nih.gov/compound/Methyl-3_4-dihydroxybenzoate). The calculated binding energy for this interaction is -6.15 kcal/mol. HE9 binds at the surface of PLpro adjacent to the binding cavity with ligands HBA and YRL. The interaction to PLpro is formed by hydrogen bonds of the dihydroxyphenol edge to the side chain of Glu 70. Hydrophobic interactions are observed including the π- stacking with the imidazole of His 73 and contacts of the benzene core to the side chain of Phe69 (Figure S2C). The NMR spectra of all the three natural compounds are presented in the supplementary information.

The described interaction networks of the three compounds involve amino acid residues Phe 69, Glu 70 and His 73 that have been previously shown to interact with ISG15 and Lys48di-Ub molecules^15, 16^. Crystal structures of SARS-CoV PLpro, in complex with Lys48di-Ub (5E6J), and SARS-CoV-2 PLpro in complex with mouse-ISG15 (6YVA) supported by molecular dynamics simulations clearly reveal the hydrophobic interactions between these amino acid residues in PLpro with either ISG15 or Lys48-di ubiquitin molecules^16^. A superimposition of the PLpro+inhibitor complex structures with PLpro+ISG15 complex (Figure 2A), shows that the binding of the natural compounds clearly disrupts and prevents the binding of ISG15 to PLpro. Critical residues Ser 22, Met 23 and Glu 27 located in the binding surface of ISG15 are no longer available to form interactions with PLpro (Figure 2B) upon binding of these natural products.

**Figure 2:**
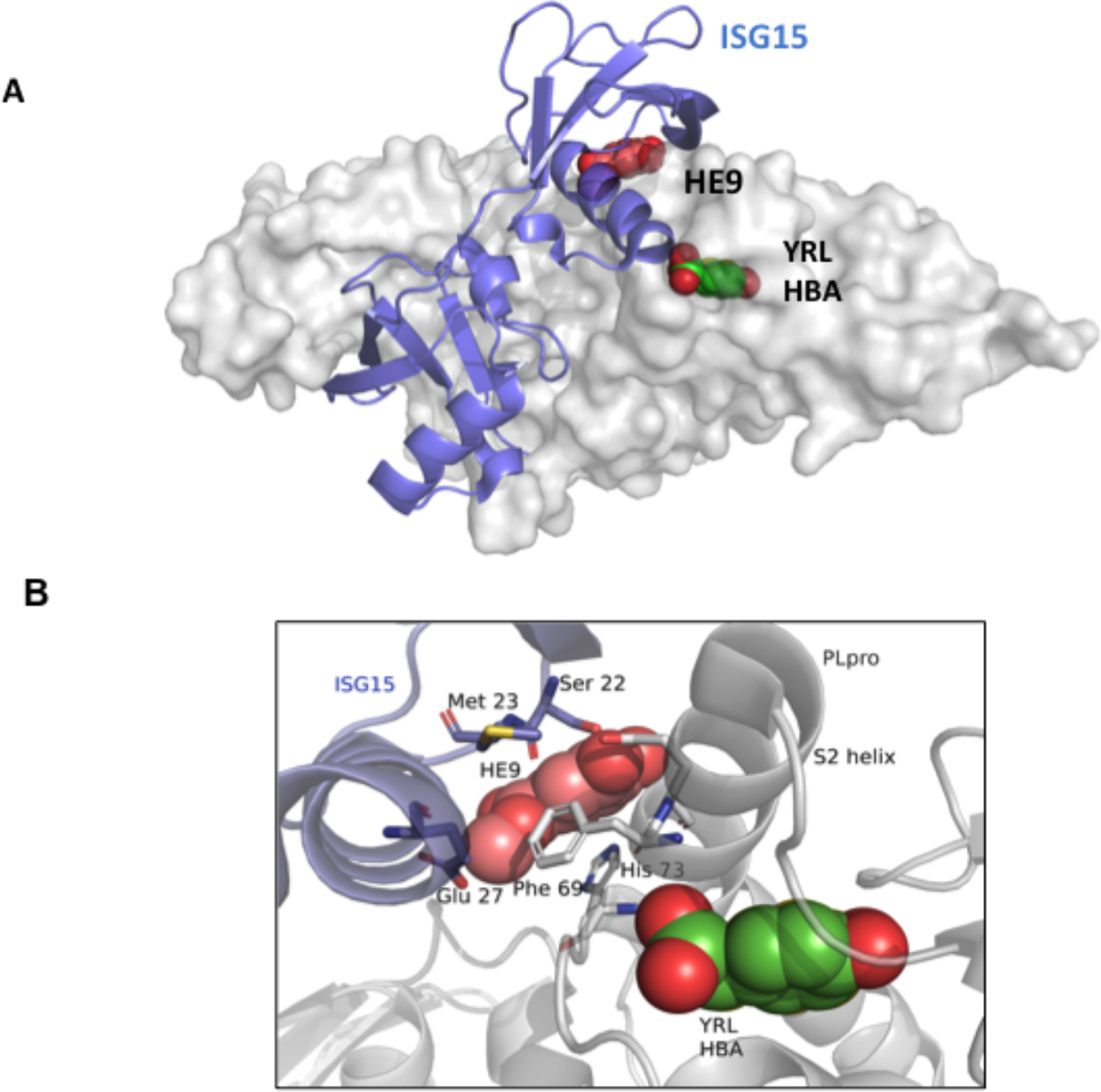
Interaction of the ISG15 molecule to PLpro is disrupted by the binding of the three natural compounds. A. Superposition of the crystal structures of SARS-CoV-2 PLpro-C111S in complex with mouse-ISG15 (PDB code 6YVA, ISG15 molecule in blue) with SARS- CoV-2 PLpro+HE9 (PDB code 7OFU, in grey surface representation). The three compounds YRL, HBA and HE9 are depicted as spheres. B. Close-up view of the ISG15 binding site. ISG15 molecule is shown as a cartoon representation (blue) with the interacting residues Ser 22, Met 23 and Glu 27 in sticks. The bound inhibitor compounds (spheres) clearly prevent the binding of the ISG15 molecule to the S2 binding site of PLpro.

PLpro enzymes share the same core residue, SARS-CoV Phe 70 and SARS-CoV-2 Phe 69 at the ISG15 binding site. A mutation of this residue in PLpro to alanine decreased the enzymatic activity, and also resulted in a slower reaction with ISG15, as compared to the wild type enzyme^16^. In MERS-CoV PLpro, Phe 69 is replaced by a lysine residue (F69K) and His 73 by a glycine residue (H73G). These variations might account for the different substrate preferences among SARS-CoV and SARS-CoV-2 and MERS PLpro. It can be seen from a superimposition of the crystal structure of the MERS-CoV PLpro+ISG15 (6BI8), with the SARS-CoV-2 PLpro-HE9 complex structure (7OFU), that F69K and H73G substitutions confer different surface properties for the interaction with ISG15 (Figures S5 A, B, C).

Cleavage of polyubiquitin chains by SARS-CoV-2 PLpro is significantly enhanced when a longer ubiquitin chain is used. This demonstrates that either Ub or ISG15 molecules bind not only to the Ub-S1 binding site but also to the Ub-S2 site, facilitated by the conserved S2 helix in PLpro, being important for the enzymatic activity^34^. Superimposition of crystal structures of PLpro in complex with Lys48 linked di-ubiquitin (5E6J) with the PLpro-HE9 complex (7OFU) showed that key residues involved in ubiquitination, Lys 11 and Lys 48 in the S2-Ub binding site, are no longer available for binding either to ubiquitin or ISG15 (Figure S4A, B). Hence, a clear molecular basis for the inhibition emerges from the three PLpro inhibitor complex structures, showing that PLpro-ISG15 interactions are affected upon the binding of the three phenolic natural products.

### *In vitro* and cell culture assays to monitor the inhibition of SARS-CoV-2 PLpro

A fluorescence deubiquitinating activity assay was carried out to assess the inhibitory effect of the three compounds (HE9, YRL, and HBA), co-crystallized with the wild type SARS-CoV-2 PLpro. A catalytically inactive PLpro mutant (C111S) was used as a control with ISG15-Rhodamine as the substrate (see materials and methods). The three natural products clearly inhibited the enzymatic reaction of PLpro. In particular the two compounds, HBA and YRL significantly decreased PLpro activity by 73%, followed by HE9 inhibiting to 55% in a deISGylation assay (Figure S6 & Table 2). It can be rationalized that the binding of the natural compounds at the S2 helix region is determinant and clearly disrupts the essential PLpro/ISG15 molecular interactions required for the enzymatic mechanism, as seen in the crystal structures of PLpro inhibitor complexes.

**Table 2.**
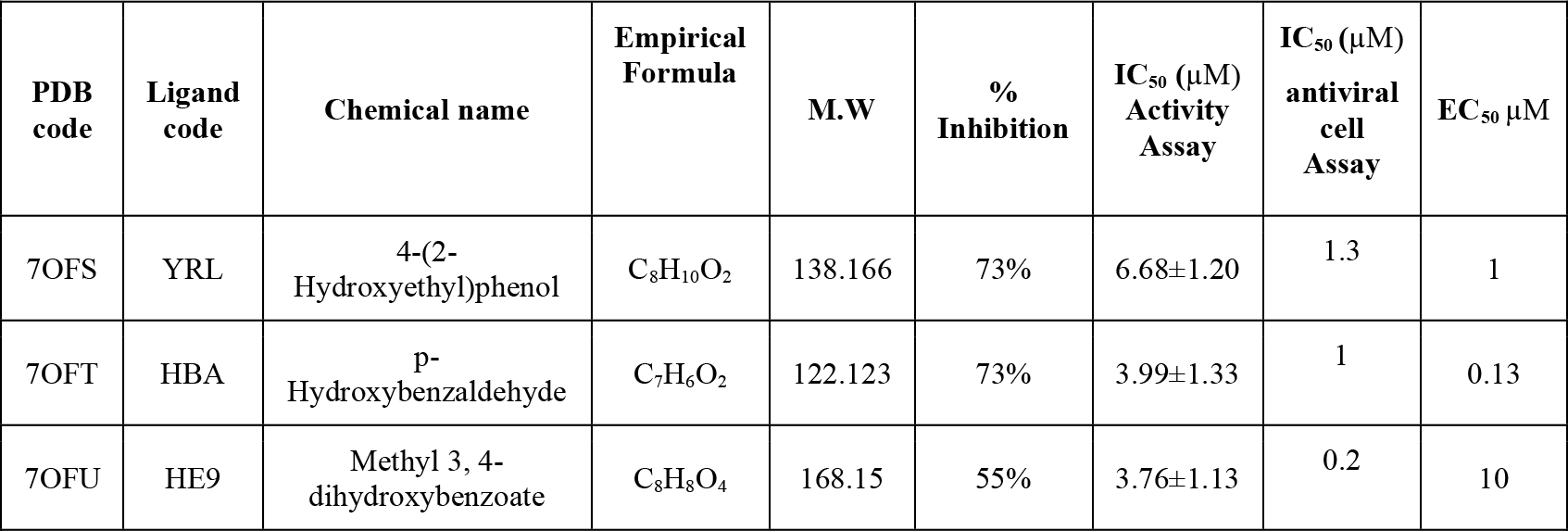
Summary of the inhibition profiles for the three PLpro inhibitors obtained from enzyme activity assays and cell line antiviral assays

Further, we determined the efficacy of inhibition of the three compounds by *in vitro* IC_50_ assays, which demonstrated a potential effect of all three natural products in a concentration range of 3.76 up to 6.68µM. The compound, GRL0617 (5-Amino-2-methyl-N-[(1R)-1-(1-naphthalenyl) ethyl] benzamide), a known inhibitor of PLpro was used as a control (Figure 3, Table 2). Antiviral activities for these natural phenolic compounds, either in crude or purified forms previously reported from several studies^35, 36^ can now be rationally related to PLpro inhibition as we show in our activity assays, in addition to other target proteins such as Mpro, that are involved in the viral replication process.

**Figure 3:**
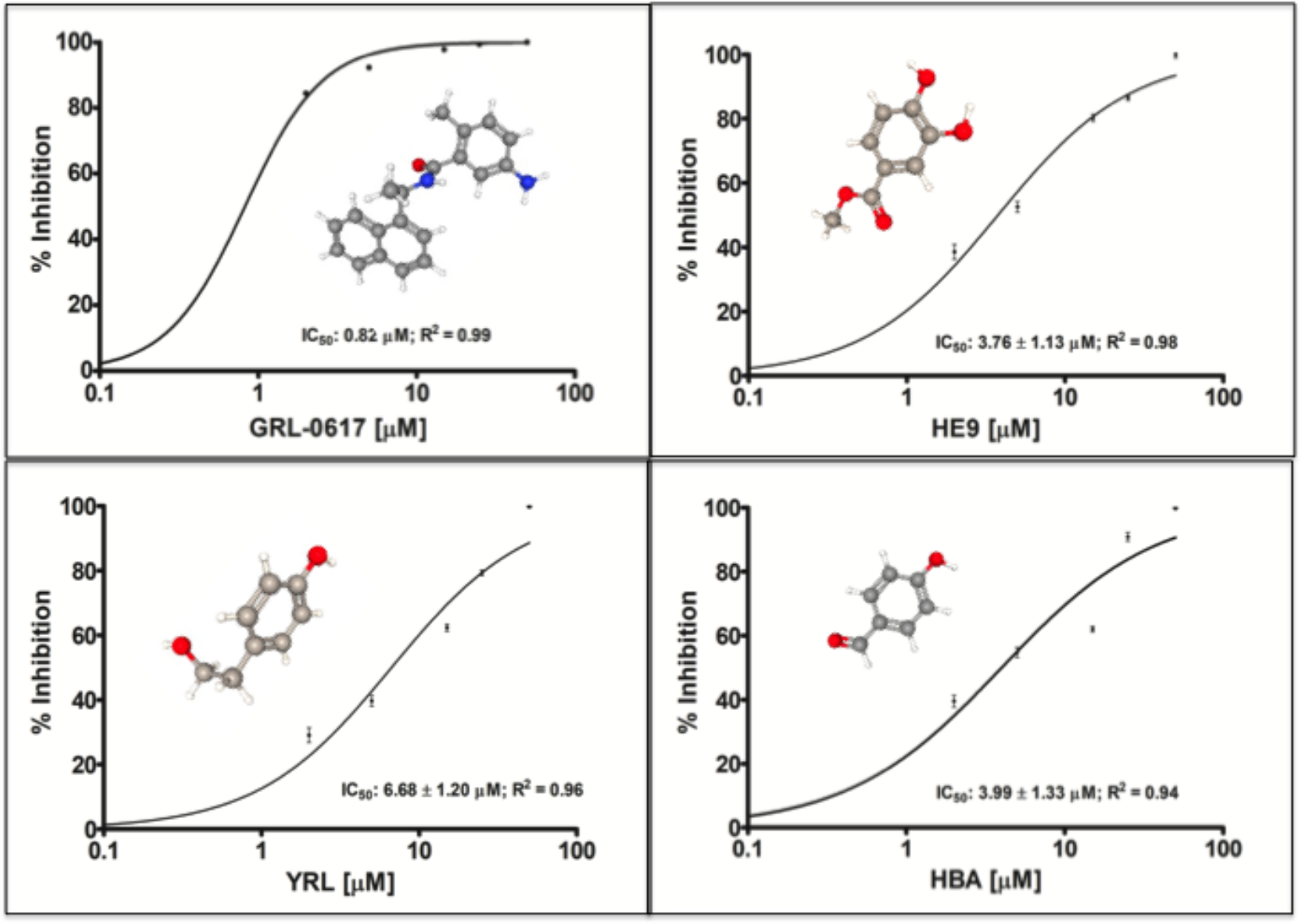
Inhibition of PLpro by the three natural compounds in deISGylation assay with ISG15- Rh substrate. IC_50_ determination was performed with ISG15-Rhodamine as the substrate at a concentration of 250nM. A gradient concentration of all three compounds YRL, HBA, HE9 and the inhibitor GRL-0617 as a control in the range from 2 µM to 50 µM was used in the reaction mixture. IC_50_ values were calculated by fitting the data to a sigmoidal dose-response-inhibition function and are presented in the log scale for interpolation. Individual data points represent the mean of normalized relative fluorescence unit per min ± SD from triplicates.

Considering the observed inhibitory synergic effect, combined with the molecular regulative antiviral homeostasis function of ISG15 in the human host, we investigated the inhibitory efficacy of the compounds HE9, HBA, and YRL towards viral replication and the cytopathic effect in living cells using Vero cell line assays.

Two distinct approaches were applied, qRT-PCR reaction as previously described^4^ and CellTiter-Glo assay, a luciferase reporter assay to determine the ATP level present in viable cells^37^. Screening experiments started at 5mM of the three compounds and used a 10-point, 1:10 dilution series with infections being performed at a multiplicity of infection (MOI) value of 0.01. The two compounds HE9 and YRL showed a reduction of viral RNA (vRNA) replication with EC_50_ values of 0.13µM and 1µM respectively, with no associated cell toxicity at 100µM (Figure 4A). Cell viability experiments were performed simultaneously under the same conditions in the absence of virus and revealed no effects on cell viability at concentrations where the compounds showed antiviral activity (Figure 4B).

**Figure 4:**
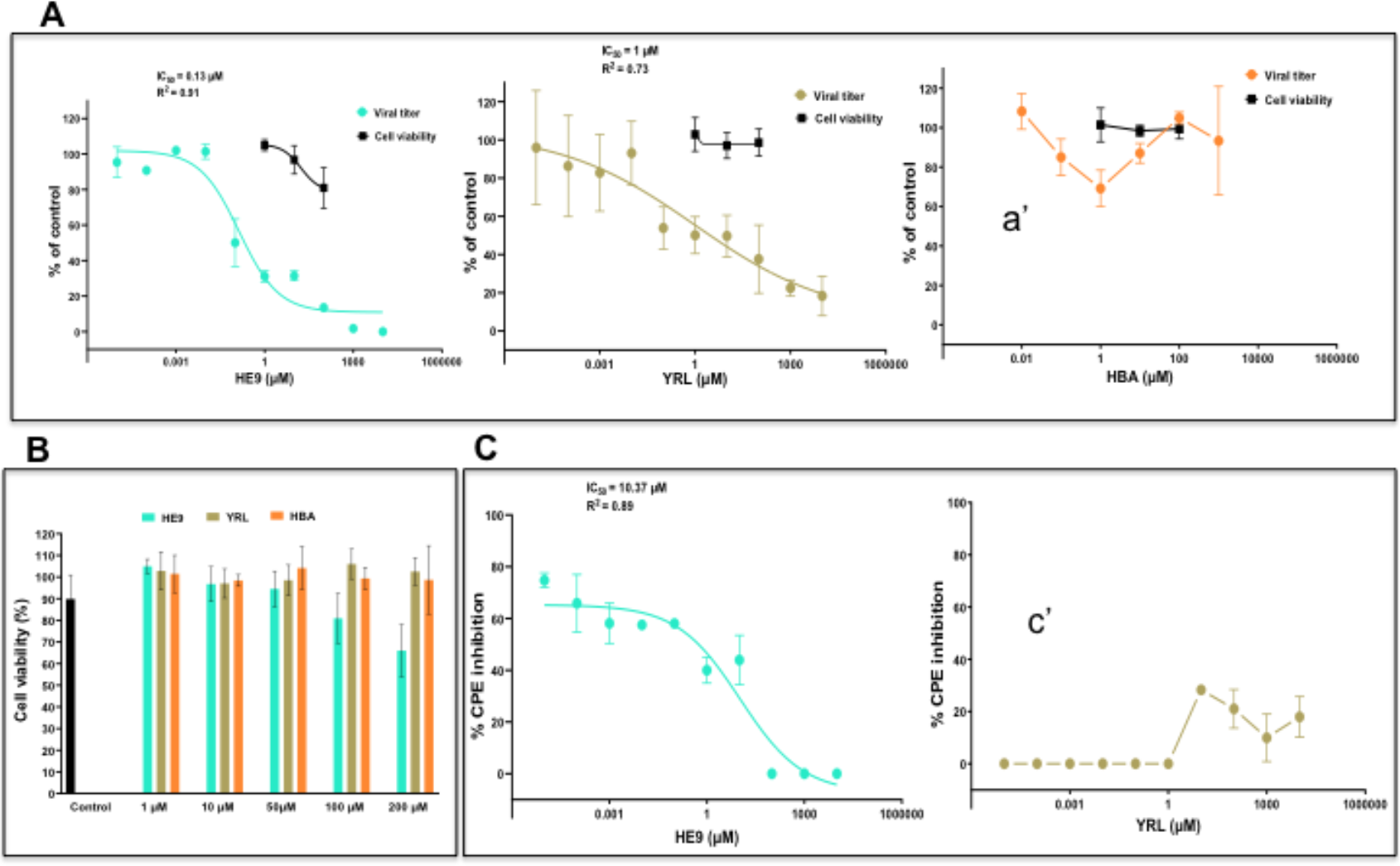
Effect of the three natural compounds on SARS-CoV-2 loading in Vero cells. A. **A.** The viral titer and cell viability were quantified by qRT-PCR (●) and CellTiter-Glo luminescence method (▪), respectively. EC_50_- and R-squared values for viral titers are shown. EC_50_-values were calculated by fitting the data to the sigmoidal function as previously described^4^. Compounds concentrations are presented in log scale for interpolation. HE9 and HBA compounds were diluted to a stock concentration of 100 mM in DMSO, while YRL was diluted in sterile water to a 50 mM stock concentration. All compounds were stored at -20°C. Individual data points represent means ± SD from four independent replicates in two biological experiments. a’. Values were plotted in a line graph with error bars displaying standard deviation. **B.** Cell viability was determined by CellTiter-Glo luminescence method. Individual data points represent means ± SD from three independent replicates in three biological experiments. **C.** CPE inhibition was determined by CellTiter-Glo luminescence method. EC_50_- and R-squared values are shown. EC_50_-values were calculated by fitting the data to the sigmoidal function. Individual data points represent means ± SD from three independent replicates in one biological experiment. c’. Values were plotted in a line graph with error bars displaying standard deviation.

The compound HE9 significantly reduced viral RNA (vRNA) replication among the three compounds studied and was further evaluated to determine the effective concentrations that can reduce not only vRNA levels but also SARS-CoV- 2 virus infectious particles applying a cytopathic effect (CPE) inhibition assay (Figure 4C). An active compound was the one which exhibited a CPE inhibition of > 50% without compromising cell viability. We were unable to fit a sigmoidal curve to the data for HBA and YRL and values are presented in a line graph (Figure 4a ’ and 4c’). Importantly, treatment of the cells with the compound HE9 reduced the viral replication and showed an ability to inhibit CPE with EC_50_ = 10µM (Figure 4C). These results from the cellular assays are in line with the *in vitro* enzymatic studies using deISGlyation assays and clearly demonstrate that the compound HE9 is a potential inhibitor of PLpro, which can protect the host cells from the viral CPE. In summary, the high resolution PLpro complex structures with phenolic natural compounds YRL, HBA and HE9 complemented by enzymatic and cellular assays, provide a molecular basis to understand the inhibitory mechanism, a route to develop effective PLpro inhibitors of ISG15 binding to PLpro and shed light on the mode of ISGylation of COVID-19 viral proteins as a new approach for preventing their interaction with human host cellular pathways. We believe that this approach to inhibit PLpro may hinder and reduce the viral ability to perform de-ISGylation in post COVID-19 viral complications, as well as providing more ISG15 within the lung tissues for the modulation of cytokine/chemokine production, to support the repair of the respiratory epithelium within COVID-19 infections^11^.

## 1. Material and Methods

### 1.1 Cloning, protein overexpression and purification of SARS-CoV-2 PLpro

A fragment of SARS-CoV-2 ORF pp1a/ab encoding the PLpro domain and corresponding to amino acids 746-1060 of non-structural protein 3 (YP_009742610.1) was cloned into pETM11(EMBL), which encodes N-terminal hexa-his tag followed by a tobacco etch virus (TEV) protease cleavage site. After cleavage by TEV protease extra two amino acids (GA) are left on the N-terminal of PLpro construct. The plasmid encoding the desired construct was transformed into *e. coli* Rosetta (DE3) cells (Merck, Germany) to perform expression via autoinduction, essentially as described before^38^ and using kanamycin for selection. An overnight cell culture was diluted and incubated in autoinduction medium containing 0.5 g L^-1^ β-D-glucose and 2 g L^-1^ lactose under constant shaking for 4 h at 37 °C and then in the presence of 100 µM ZnCl_2_ additionally over-night at 18 °C. Subsequently, cells were harvested by centrifugation and disrupted by sonication in lysis buffer (50 mM NaH_2_PO_4_, 150 mM NaCl and 10 mM imidazole, pH 7.2).

Cell extracts were maintained at 4 °C and centrifuged at 12,000 × g for 1 h. The clear supernatant was incubated with Ni-NTA affinity resin (Thermo Fisher Scientific, USA). PLpro was eluted by gravity flow using lysis buffer supplemented with 300 mM imidazole and subsequently incubated with TEV protease at a molar ratio of 20:1 in the presence of 1 mM DTT. Cleavage was performed during dialysis against 50 mM Tris, 150 mM NaCl and 1 mM DTT adjusted to pH 7.3 and for 14 h at 8 °C. After removing protease and the cleaved off tag by affinity chromatography, PLpro was purified to homogeneity using size-exclusion chromatography, i.e. a HiLoad 16/600 Superdex 75 column connected to an ÄKTA purifier (GE Healthcare, GB) equilibrated with 50 mM Tris, 150 mM NaCl and 1 mM TCEP at pH 7.5. Purity and integrity of the protein were verified via SDS polyacrylamide gel electrophoresis and DLS (Dynamic Light Scattering). The concentration of PLpro with a calculated molecular weight of 35760 Da (εcalculated = 45270 M^-1^ cm^-1^) was adjusted to 20 mg mL^-1^ in preparation for vapor diffusion crystallization trials.

### 1.2. Crystallization of ligand free SARS-CoV-2 PLpro and complexes with compounds

Initial crystallization screening experiments were performed using the sitting-drop vapour diffusion method utilizing the Oryx4 robot (Douglas Instruments) with the SWISSCI 96-well plates. Wizard™ Classic 1, 2, 3 and 4, JCSG+, PACT crystallization formulations were tried for initial screening experiments. Crystallization was performed with a protein:reservoir ratio of 2:1 at 4 °C and 20 ^°^C. Initial hits were obtained from the Wizard screen, condition G11 (0.1M acetate buffer pH 4.5, 0.8M NaH_2_PO4/1.2M K_2_HPO_4_) at 4 °C. Further optimization was done by changing the buffer to 0.1M Tris-HCl pH = 8.0 and including 10% glycerol that resulted in 0.2-0.3mm bipyramidal crystals. These crystals diffracted X-rays to a resolution of 1.42 Å.

PLpro complex crystals with compounds were grown by the co-crystallization method using the same condition as summarized above. 100 nL droplets of 10 mM compound solutions in DMSO from the Sadia library of natural compounds were applied onto a 96-well SWISSCI plate and the compounds were dried in vacuum before the addition of 200 nL of (20 mg/mL) PLpro protein solution and 100 nL of the crystallization condition (0.1 M Tris-HCl buffer pH 8.0, 0.8 M NaH_2_PO_4_/1.2 M K_2_HPO_4_ and 10% glycerol). The drops were equilibrated in a sitting drop vapour diffusion setup with 80 µl of reservoir solution. The plates were incubated at 4 °C and crystals appeared in 2 days and grew reproducibly to dimensions of approx. 0.2 × 0.3 × 0.2 mm^3^ in 4 days. Crystals were manually harvested directly from the drop and flash-frozen in liquid nitrogen for diffraction data collection.

### 1.3. Data collection, structure solution and refinement

Diffraction data from the ligand-free and complex PLpro crystals were collected at beamline P11, PETRA III / DESY, Hamburg. All datasets of ligand-free PLpro were processed using the program XDS^39^ with a reference dataset to ensure consistent indexing. From a total of 64 complete datasets, the strongest were selected based on (I/σ)^asymptotic^ greater than 20^40^. These 25 datasets were then subjected to iterative merging using CODGAS^41^ run with standard parameters. The best merged datasets was further manually filtered leading to the final dataset that contained five datasets. These were scaled with XSCALE^39^ and final merging and resolution cut-off was applied using AIMLESS^42^. Structure solution was achieved by molecular replacement method with PHASER^43^ using the PLpro coordinates with PDB code 7JRN as search model. Successive rounds of manual building with the program COOT^44^ and refinement with PHENIX^45^, the addition of phosphate, glycerol, chloride ions and water solvent molecules to the model, followed by a final round of TLS refinement completed the structure refinement at a resolution of 1.42 Å.

An automatic data processing pipeline, hit finding, clustering^46^, PanDDA analysis ^47^and refinement protocols as described previously^4^ were used for the structure solution and analysis of PLpro in complex with the natural compounds from the library consisting of 500 compounds. Data processing with XDS resulted in 1469 datasets and includes more than one dataset per compound. Data quality indicators CC_1/2_ and Wilson B-factors are plotted as shown in Figure S7. Final rounds of manual refinement with either Refmac^48^ or Phenix^49^ together with manual model building applying COOT resulted in the final refined structures. Data collection and refinement statistics for PLpro and complexes are summarized in Table 1, supplementary information. All figures were prepared using PyMol^50^.

### 1.4. SARS-CoV-2 PLpro inhibition assays and IC_50_ determination

Activity assays were performed for SARS-CoV-2 PLpro native and mutant enzyme (PLpro C111S mutant) to determine the deISGylation activity effected by the three natural compounds, following previously published protocols^16, 17, 51^. The assays were performed with a total reaction volume of 100 µL in non-binding, black bottom, 96-well plate and reactions were measured on a Tecan Infinite M plus plate reader (Tecan Group Ltd, Switzerland) using optical settings specific for ISG15- Rhodamine (UbiQ-127, UbiQ Bio). ISG15-Rhodamine is a fluorogenic substrate that contains the cleavage sequence RLRGG recognized by PLpro at the C-terminus. The cleavage of the amide bond between the terminal glycine residue and the rhodamine110 fluorophore releases the fluorescent Rh110-morpholinecarbonyl that results in an increase of fluorescence intensity, measured as RFU (Relative Fluorescence Unit). The fluorophore has an excitation and emission at 492nm and 525nm respectively. The ISG15 substrate (UbiQ-127) and PLpro were used at a final concentration of 100nM and 10nM respectively. Relevant substrate and positive controls (GRL0617) was used throughout the assay. SARS-CoV-2 PLpro native, mutant and substrates were diluted in assay buffer (20 mM Tris-HCl pH 7.5, 150 mM NaCl, 1 mM TCEP) and reactions were started upon addition of PLpro in a final volume of 100 µL and measured at 25 °C. The putative inhibitor compounds were incubated with PLpro enzyme at 10 °C for six hours. Inhibition kinetics were measured in triplicates over 60 minutes with one read per minute in two independent experiments. Measured fluorescence values were blank corrected with buffer containing the ISG15-Rhodamine substrate.

IC_50_ determination was performed with ISG15-Rhodamine as the substrate at a concentration of 250nM. The assays were performed as described above, however a gradient concentration of all three natural compounds and GRL-0617 were used in the concentration ranging from 2 µM to 50 µM in reaction mixture prior to incubation. The IC_50_ values were calculated by the dose-response-inhibition function after the normalization of the enzymatic activity values. Microsoft Excel and GraphPad Prism (version 8.3.1) were used for analyzing the results and preparation of corresponding figures.

### 1.5 Cytotoxicity assays

Vero cell lines (ATCC® CCL-81™) were cultivated in Dulbecco’s modified Eagle’s medium (DMEM) supplemented with 10% fetal bovine serum (FBS). The cells were seeded in 96-well plates at a density of 3.5×10^4^ cells/well, following 24 hours incubation at 37°C and 5% CO_2_ atmosphere. The cell culture media was changed and ten-fold serial dilutions of the compounds were added. Cell viability following 72 h treatment of cells with the respective compounds was determined via CellTiter-Glo® Luminescent Cell Viability Assay (Promega), following manufacturer’s instructions. Luminescent signal was recorded using a CLARIOstar multi-mode microplate reader (BMG Labtech, Germany). Data were obtained from three independent replicates in three biological experiments. Samples deemed to be technical failures and extreme outlier were removed. Wells containing only culture medium served as a control to determine the assay background.

### 1.6 Antiviral activity assay

Vero cell lines (ATCC® CCL-81™) cultivated in DMEM supplemented with 10% FBS was seeded in 96-well plates at a density of 3.5×10^4^ cells/well, following 24 hours incubation at 37°C and 5% CO_2_ atmosphere. The cell culture media was changed and tenfold serial dilution of the compounds were added to the cells. The assays were performed as published previously^4^. Briefly, after 1 h incubation, SARS-CoV-2 strain^52^, diluted in DMEM with 2.5% FBS, was added to the cells at a MOI of 0.01 and allowed absorption for 1 h. The viral inoculum was removed, and cells were gently washed with phosphate-buffered saline (PBS) without calcium and magnesium. Fresh DMEM with 2.5% FBS containing the compounds was added back onto the cells. Cell culture supernatant was harvest 42 h post-infection and viral RNA was purified using MagMAX™ Viral/Pathogen Nucleic Acid Isolation Kit (Thermo Fisher Scientific). The samples were processed using the semi-automated NucliSENS® easyMag® platform (bioMérieux, Lyon, France), following the manufacturer’s’ instructions. All SARS-CoV-2 infections were performed in a biosafety level 3 laboratory at the Institute of Biomedical Sciences, University of São Paulo, Brazil. The viral titers were determined by the qRT-PCR method using AgPath-ID™ One-Step RT-PCR Kit (Thermo Fisher Scientific) and a sequence of primers and probe for the E gene^53^. The viral titers were calculated using a standard curve generated with serial dilutions of a template known concentration and expressed in TCID_50_/mL. Infected cells with the addition of 0.5% DMSO were used as control. EC_50_-values were calculated by fitting the data using GraphPad Prism version 8.00 (GraphPad Software, La Jolla California USA). Data were obtained from four independent replicates in two biological experiments. Samples deemed to be technical failures and extreme outlier were removed.

### 1.7 Cytopathic effect inhibition

Vero cell lines (ATCC® CCL-81™) cultivated in DMEM supplemented with 10% FBS were seeded in 96-well plates at a density of 3.5×10^4^ cells/well, following 24 hours incubation at 37°C and 5% CO_2_ atmosphere. The cell culture media was changed and tenfold serial dilution of the compounds were added to the cells. The cells were infected at MOI 0.01 and the cytopathic effect (CPE) inhibition following 42 h treatment of cells with the respective compounds was determined via CellTiter-Glo® Luminescent Cell Viability Assay (Promega). Luminescent signal was recorded using a CLARIOstar multi-mode microplate reader (BMG Labtech, Germany). Data were obtained from three independent replicates in one biological experiment. Samples deemed to be technical failures and extreme outlier were removed.

The luminescent-based assay measures the inhibition of SARS-CoV2–induced cytopathic effect (CPE) in Vero cell line (ATCC® CCL-81™)^37^. Percent cytopathic effect (CPE) inhibition was defined as [(test compound - virus control)/(cell control - virus control)] *100. EC_50_ values were fitted by sigmoidal function using GraphPad Prism version 8.00 (GraphPad Software, La Jolla California USA).

## Data availability

Coordinates and structure factors were deposited in the Protein Data Bank PDB, with codes: 7NFV (PLpro), 7OFS (PLpro in complex with YRL, 4-(2- hydroxyethyl)phenol), 7OFT (PLpro in complex with HBA, p-hydroxybenzaldehyde) and 7OFU (PLpro in complex with HE9, 3, 4- dihydroxybenzoic acid, methyl ester).

## ACKNOWLEDGEMENTS

We acknowledge Deutsches Elektronen-Synchrotron (DESY, Hamburg, Germany), a member of the Helmholtz Association HGF, for the provision of experimental facilities, the Federal Ministry of Education and Research (BMBF) *via* projects 05K2020, 05K19GU4, the Joachim-Herz-Stiftung Hamburg (project Infecto- Physics), and financial support *via* the collaborative project between Universities São Paulo (USP) and Hamburg, (UHH) UHH-USP-FAPESP Sprint Project 2019. The authors also acknowledge the support of the Cluster of Excellence ’Advanced Imaging of Matter’ of the Deutsche Forschungsgemeinschaft (DFG) - EXC 2056 - project ID 390715994. We also acknowledge the contributions of ICCBS (International Center for Chemical and Biological Sciences) collaborators, Prof. Dr. M. Iqbal Choudhary, Prof. Dr. Bina S. Siddiqui, Prof. Dr. Shaiq Ali, Prof. Dr. Sabira Begum and Prof. Dr. Atia-tul-Wahab for providing the library of compounds mentioned in this manuscript.

## COMPETING INTERESTS

The authors declare no competing interests.

## ADDITIONAL INFORMATION

Correspondence and requests for materials should be addressed to.

## Supplementary Materials

This PDF file includes:

Supplementary Text Figs. S1 to S8 Tables S1 to S2

### Supplementary Text

#### I. Extraction, Isolation and Purification of p-hydroxy-benzaldehyde (HBA)

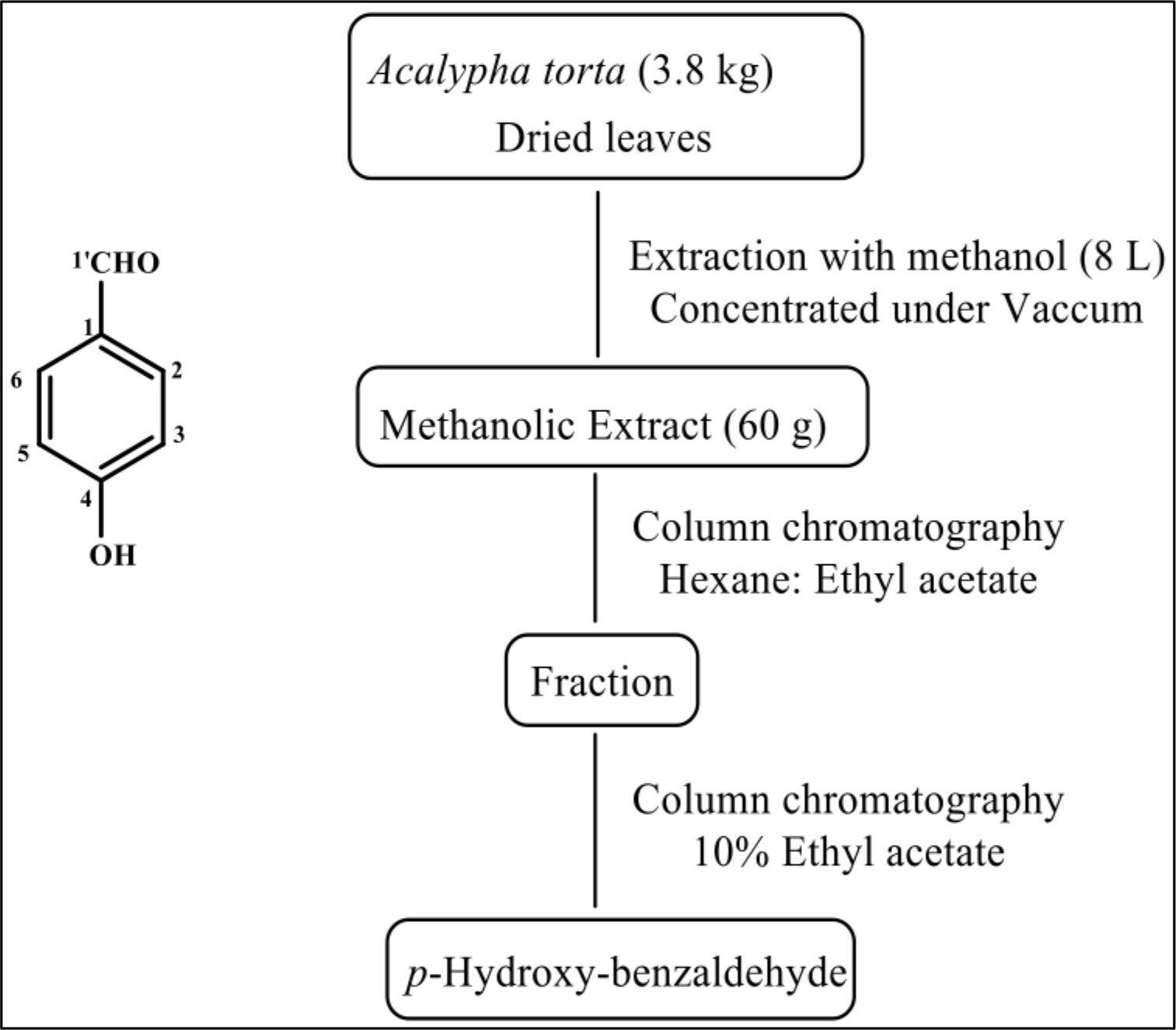

Leaves of *Acalypha torta* were collected from the flower Kingdom in Ibada, Oyo State, Nigeria. The compound p-hydroxy-benzaldehyde (HBA) was extracted, isolated and purified following the scheme above. The compound was analyzed by Electron Ionization Mass Spectrometry (EI-MS) that showed a peak with m/z 122, with a fragmentation pattern at m/z 121, 97, 95, 81, 69, and 55. The NMR spectrum is shown below. The MS and NMR spectra of HBA corresponds to the data published before^54^.

NMR spectra of p-Hydroxy-benzaldehyde (HBA)

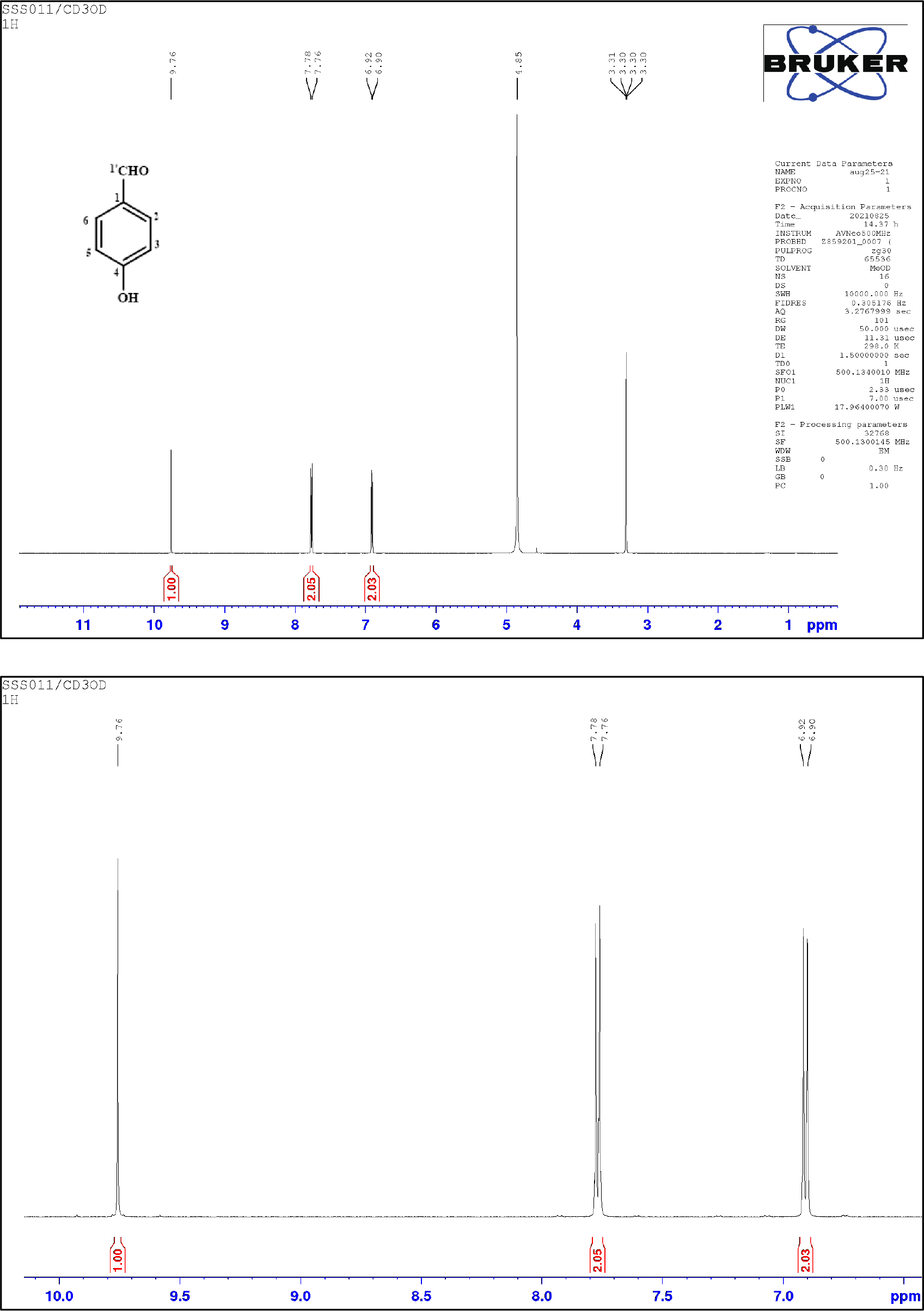

#### II. Extraction, Isolation and Purification of 4-(2-Hydroxyethyl) Phenol (YRL)

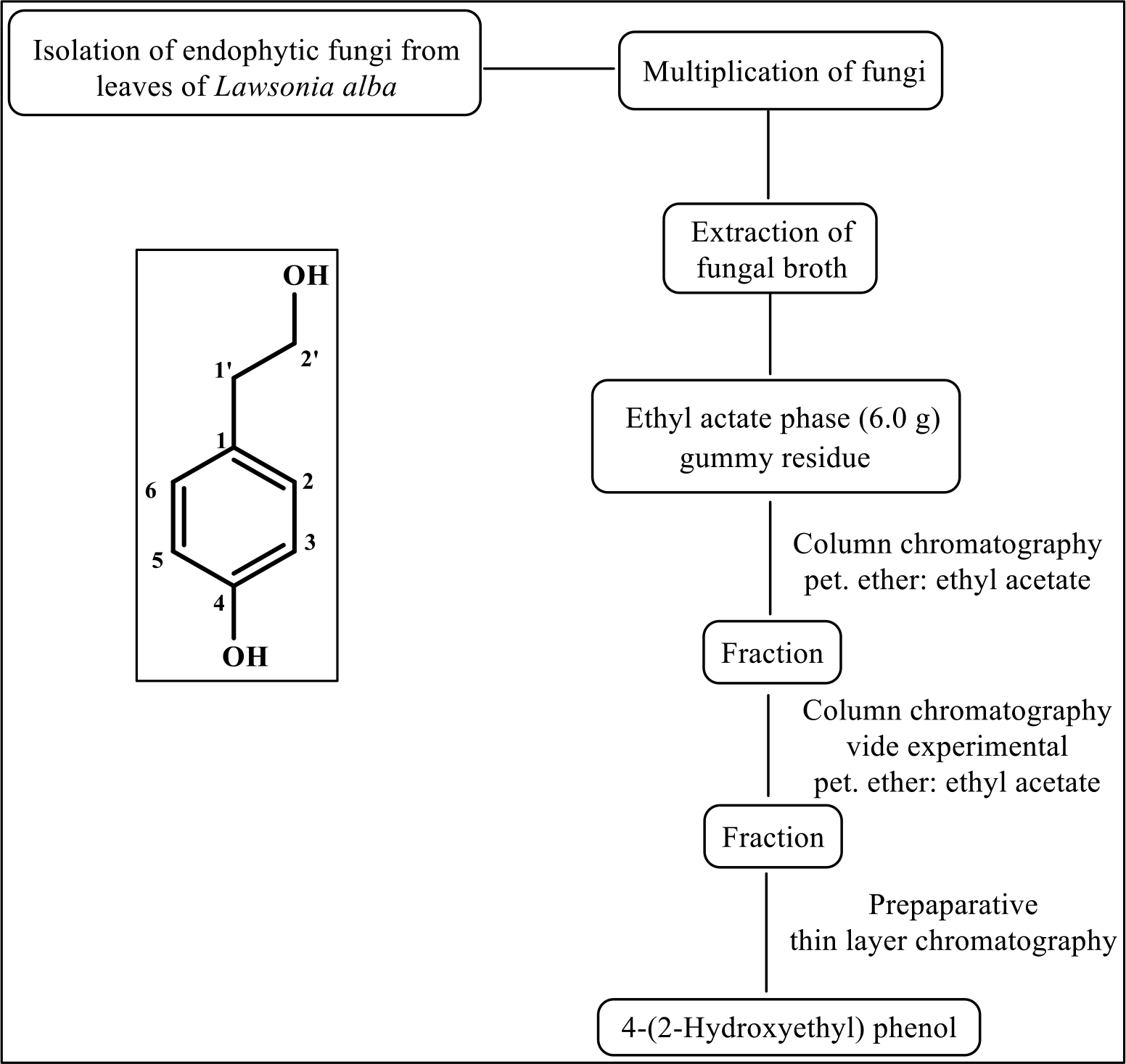

Endophytic fungi from leaves of *Lawsonia alba* were collected at ICCBS, University of Karachi, Karachi, Pakistan. Extraction, isolation and purification of 4-(2-hydroxyethyl) phenol (YRL) is shown in the scheme above. The compound YRL was analyzed by mass spectrometry that showed a peak at m/z corresponding to a mass of 138.07 Da. The NMR spectrum is shown below. Spectral data obtained for the compound were identical to previously reported data^55^.

NMR spectra of 4-(2-Hydroxyethyl) Phenol (YRL)

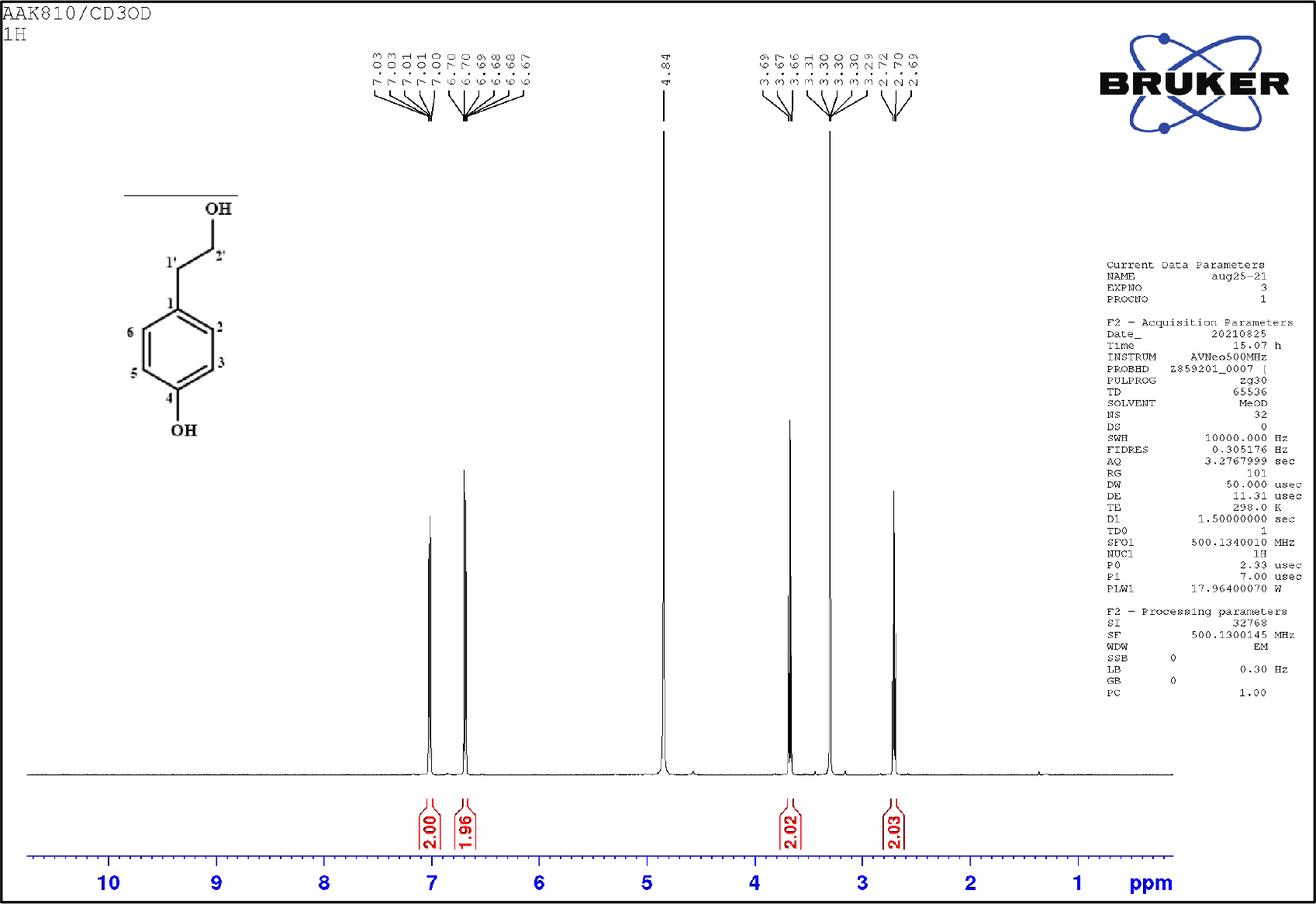
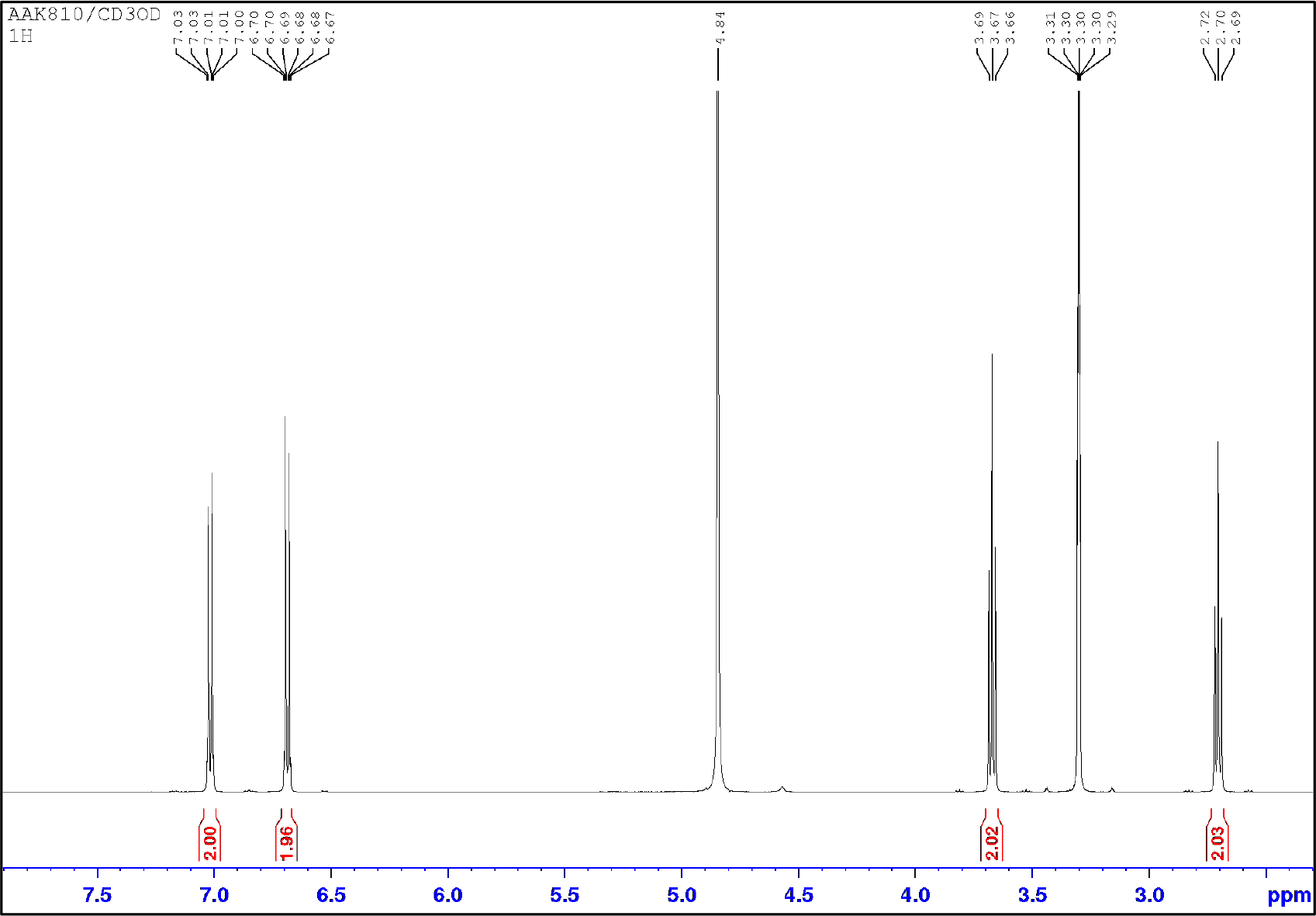

#### III. Extraction, Isolation and Purification of Methyl-3,4-dihydroxybenzoate (HE9)

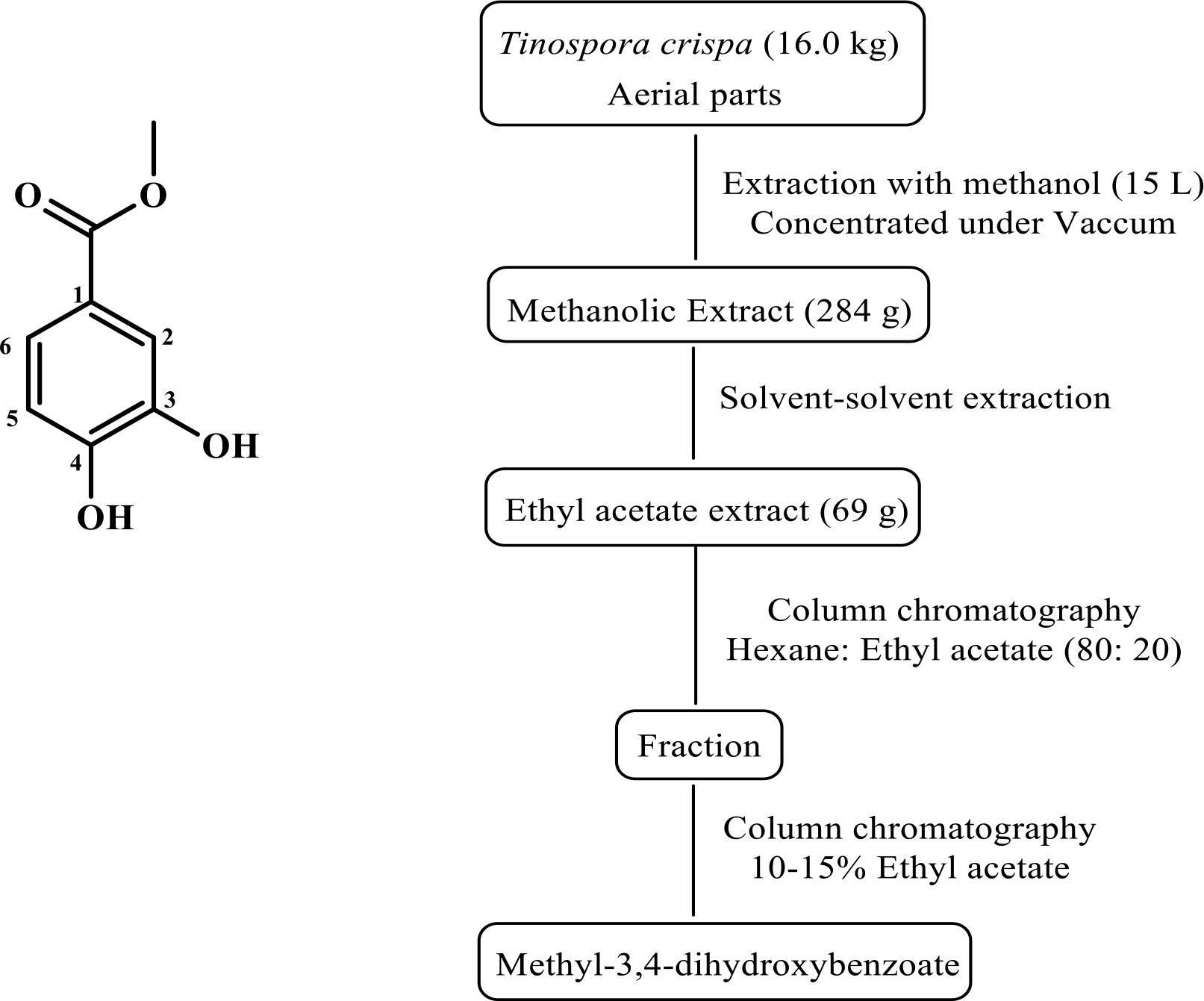

Aerial parts of *Tinospora crispa Meirs* were collected from the herbal gardens of the Laboratory of Natural Products, University of Putra, Malaysia. Extraction, isolation and purification of methyl-3,4-dihydroxybenzoate (HE9) is shown in the scheme above. The compound HE9 was analyzed by mass spectrometry that showed a peak at m/z corresponding to 168.04 Da. The NMR spectrum is shown below. The spectra are in agreement to previously reported data of the same compound isolated from *Schisandra verruculosa*^56^.

NMR spectra of Methyl-3,4-dihydroxybenzoate (HE9)

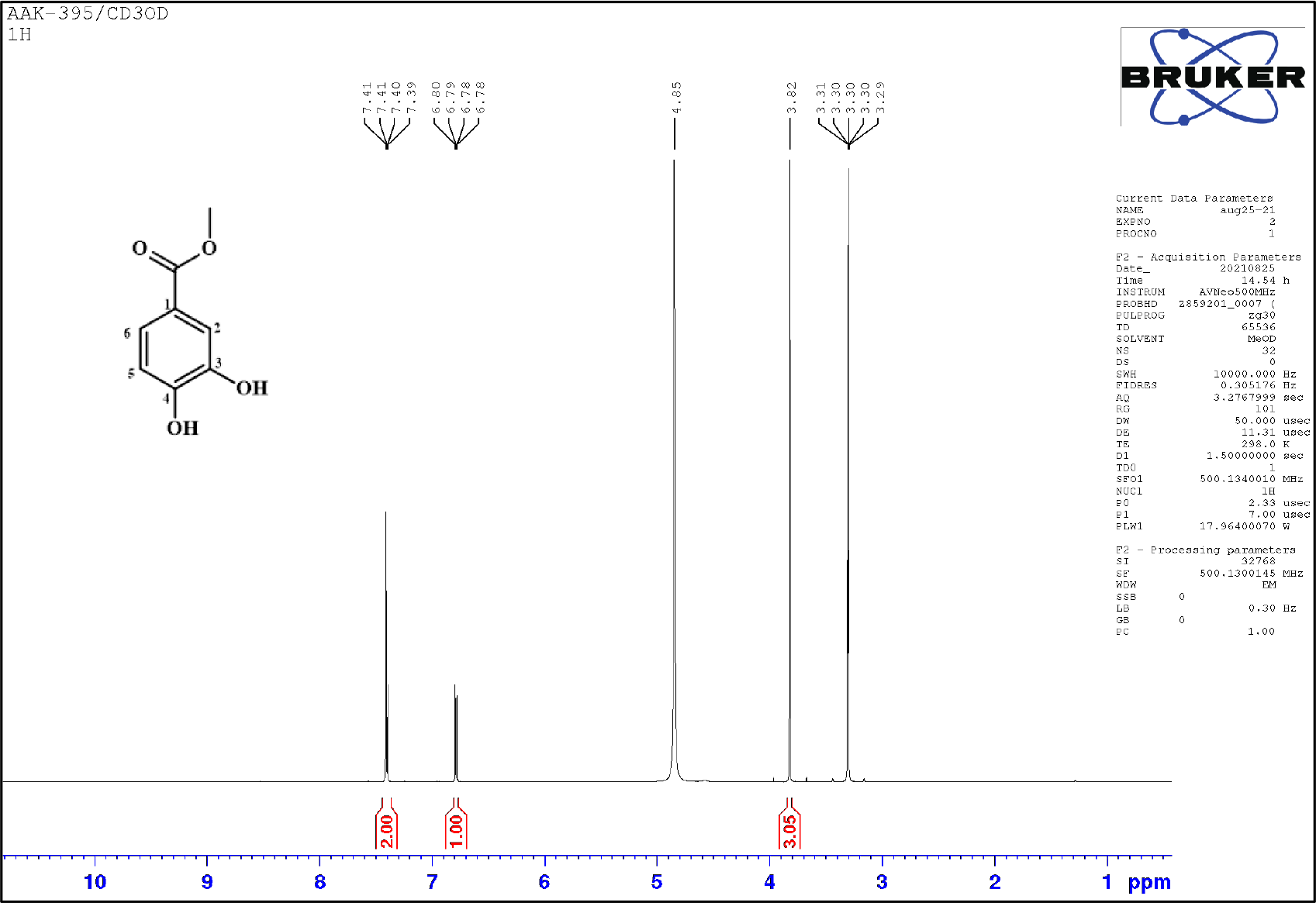
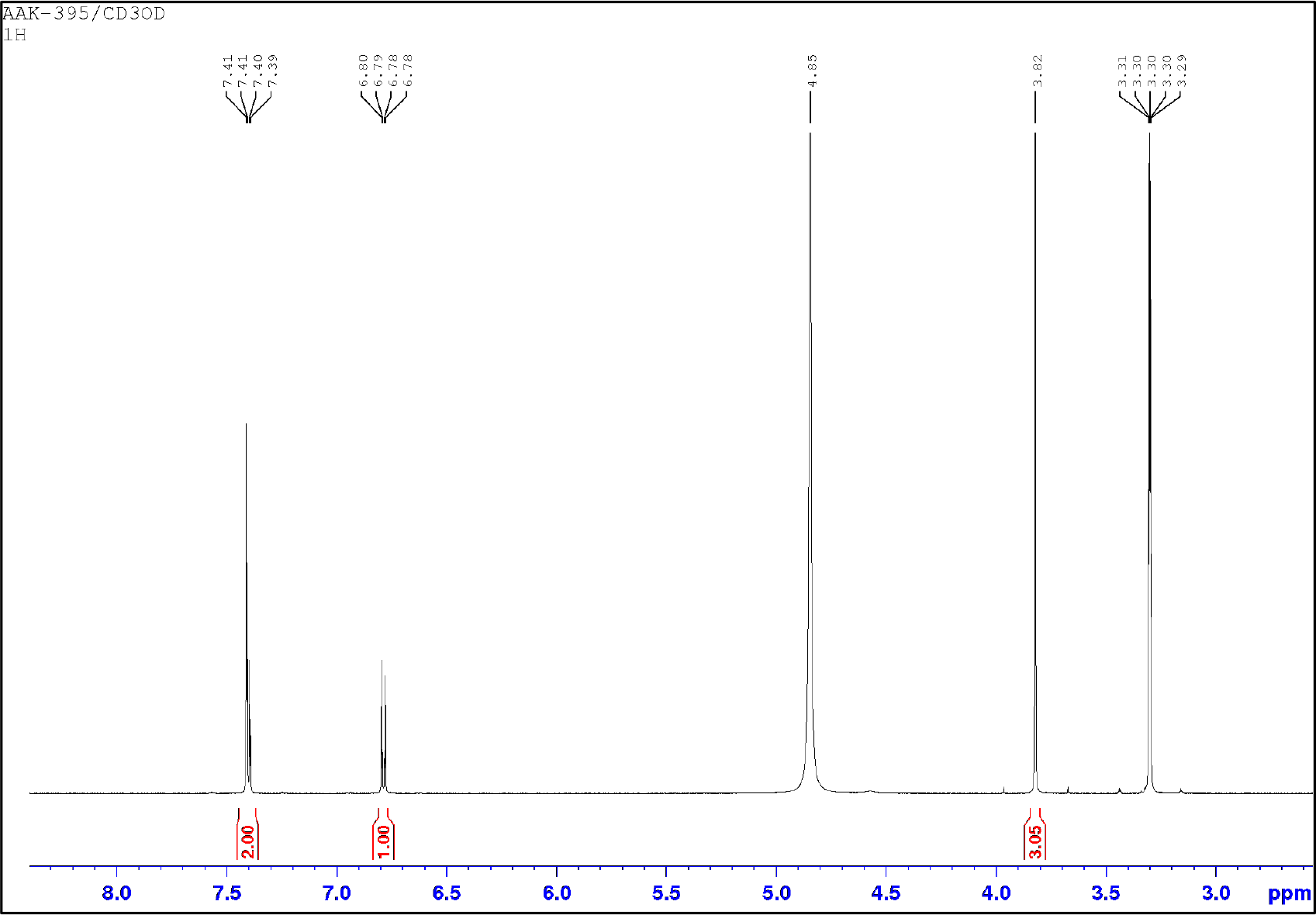

**Fig. S1.**
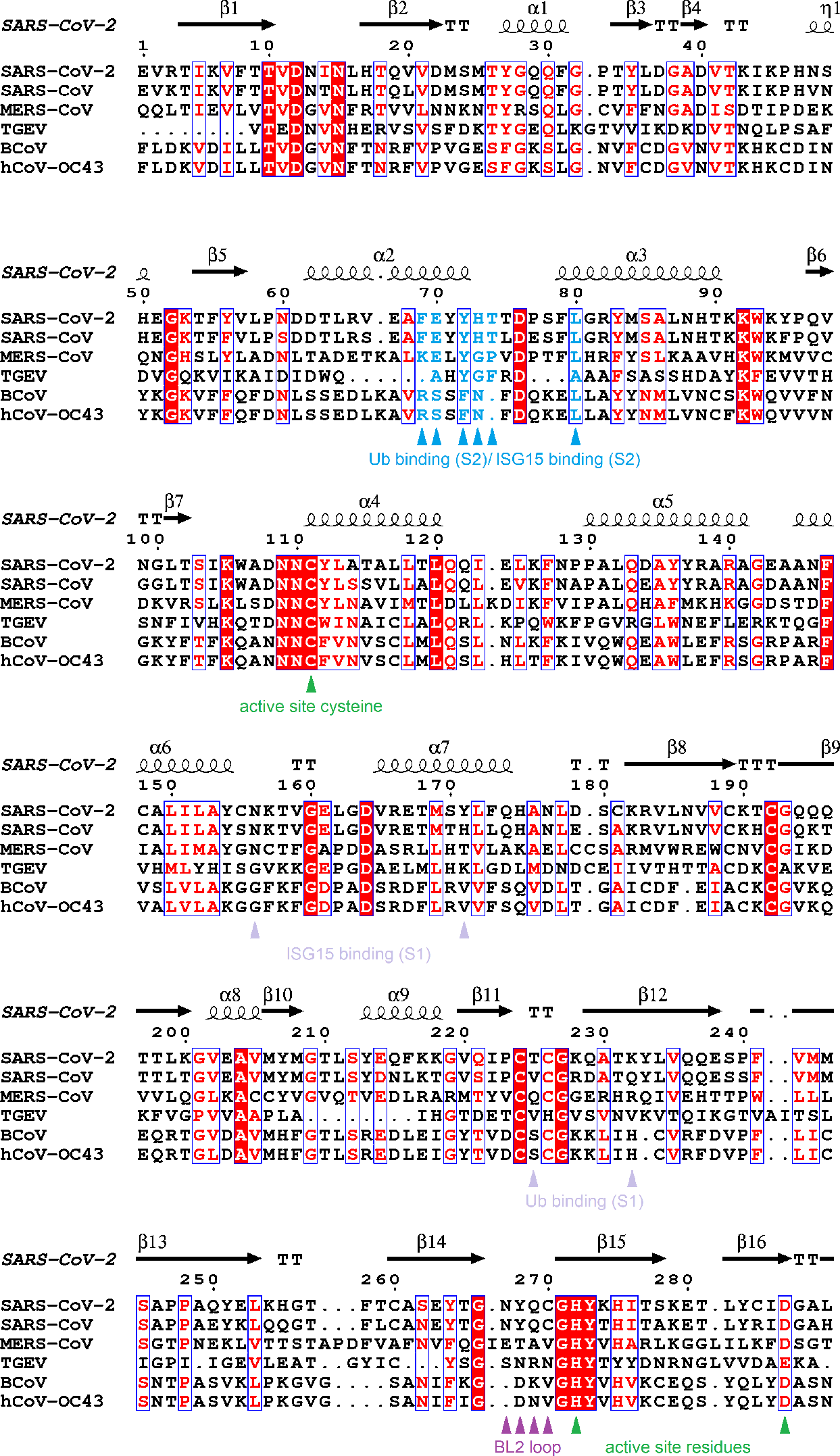
Sequence alignment of PLpro domains from selected α-, β- and related γ- coronaviruses. Sequences of PLpro from SARS-CoV-2, ID: P0DTC1; SARS-CoV, ID: P0C6X7; MERS-CoV (Middle East Respiratory Syndrome Coronavirus), ID: K9N638; TGEV (Transmissible Gastroenteritis Virus), ID: P0C6V2; BCoV (Bovine Coronavirus), ID: P0C6W7; hCoV-OC43 (Human Coronavirus OC43), ID: P0C6X6 are aligned using Espript 3^57^. The secondary structure and sequence numbering based on our crystal structure of PLpro (PDB code 7NFV) is depicted on the top of the alignment. Active site residues are indicated in green triangles, the blocking loop in purple, the ISG15 binding S1 site in dark red and the ISG15 S2 allosteric binding site overlapping with the inhibitor binding site is shown in light blue.

**Fig. S2A.**
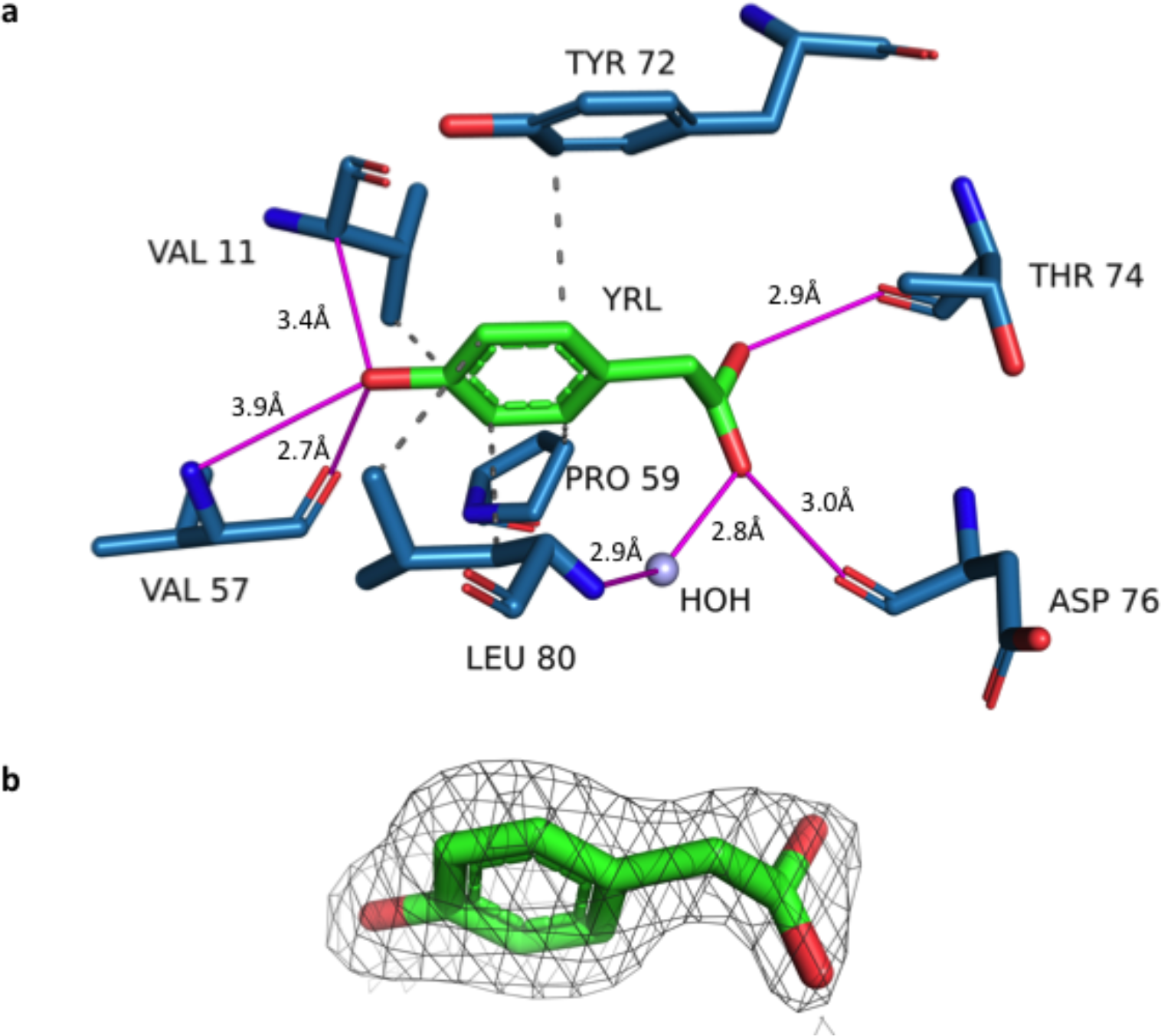
Interaction network of the three compounds in PLpro complex structures. A. YRL a. YRL interactions with PLpro are mediated by hydrogen bonds of the phenolic hydroxyl to amino acid residues Val 11 and Val 57. The aliphatic hydroxyl substituent of YRL has an alternate conformation with equal occupancy in the complex. Both conformations are stabilized by hydrogen bonds to backbone carbonyl oxygens of Thr 74 and Asp 76. Hydrophobic interactions (black dashed line) with Val 11, Pro 59, Tyr 72 and Leu 80 complete the interaction network. YRL is represented as green sticks and the amino acid residues in blue sticks. b. The 2Fo-Fc electron density map contoured at a sigma level 1.0 for YRL

**Fig. S2B.**
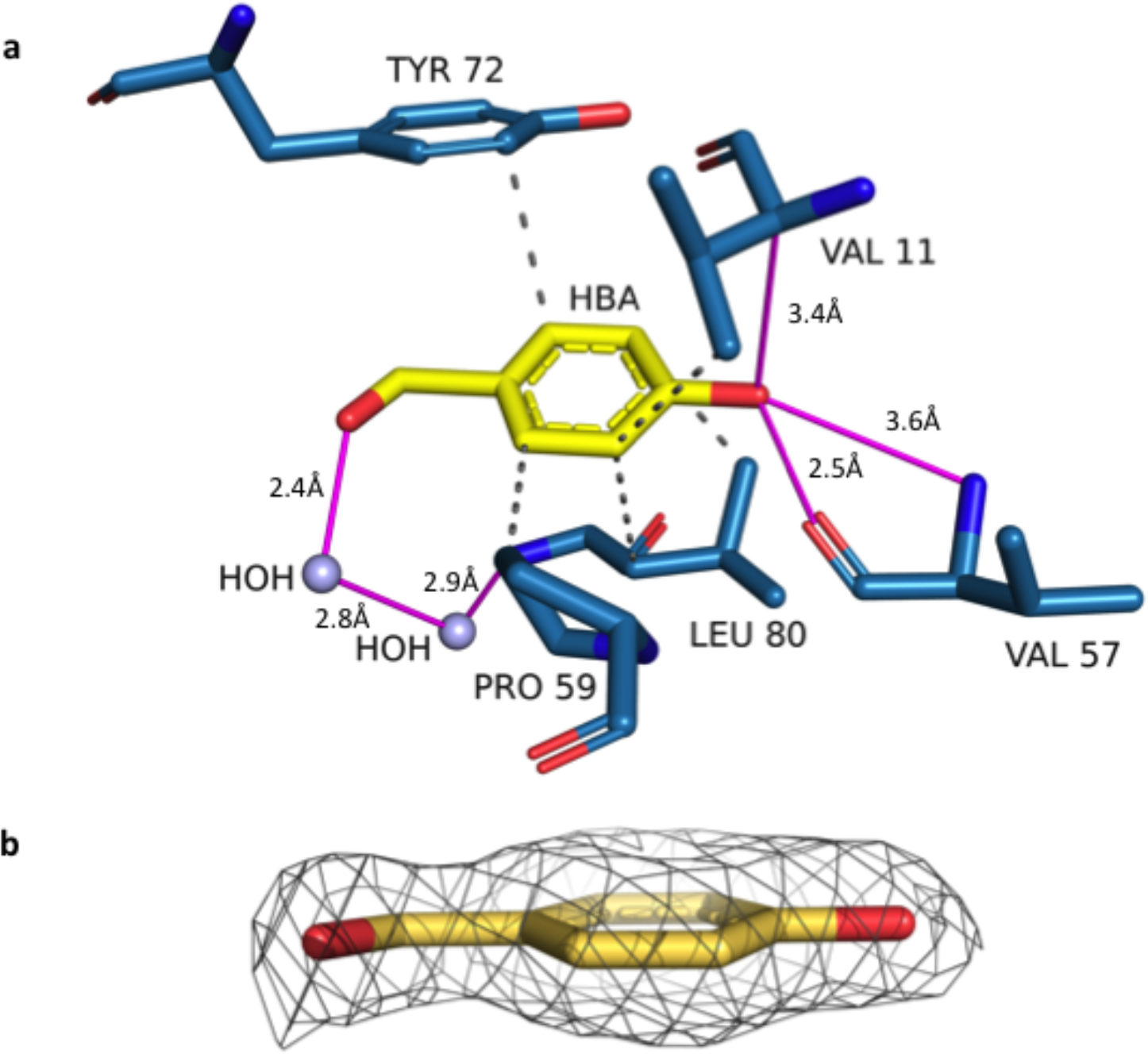
Interaction network of the three compounds in PLpro complex structures. a. HBA binds in a hydrophobic cavity with interactions to Val 11, Pro 59, Tyr 72 and Leu 80. The binding pattern is similar to that of YRL, including the hydrogen bond to Val 57. HBA is represented as yellow sticks and the amino acid residues in blue sticks. b. The 2Fo-Fc electron density map contoured at a sigma level 1.0 for HBA

**Fig. S2C.**
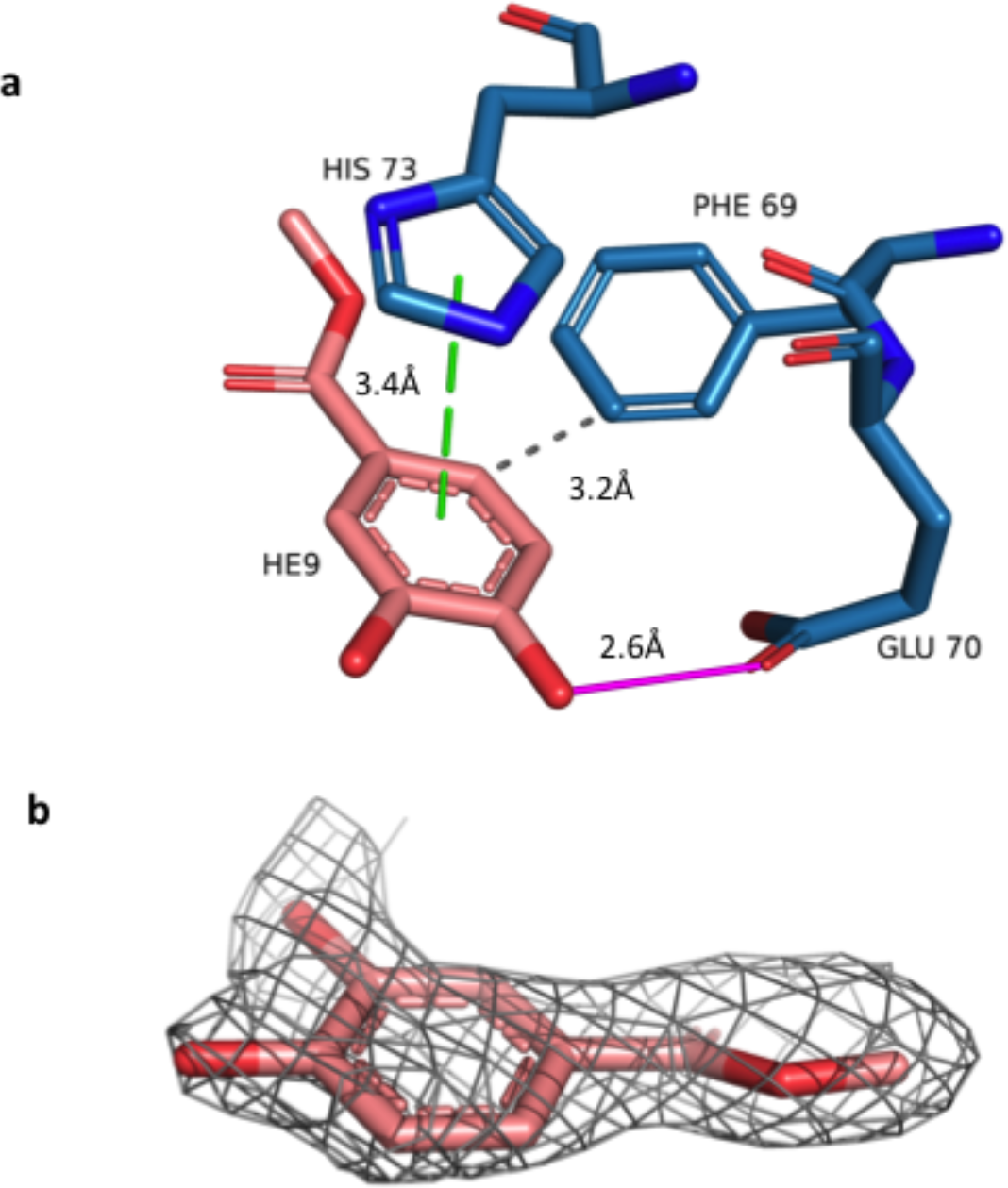
Interaction network of the three compounds in PLpro complex structures. a. HE9 binding to PLpro is mediated by hydrophobic interaction with Phe 69 and π- stacking (green dashed line) with His 73. A hydrogen bond with the side chain of Glu 70 completes the interaction network of HE9 to PLpro. HE9 is represented as pink sticks and the amino acid residues in blue sticks. b. The 2Fo-Fc electron density map contoured at a sigma level 1.0 for HE9

**Fig. S3.**
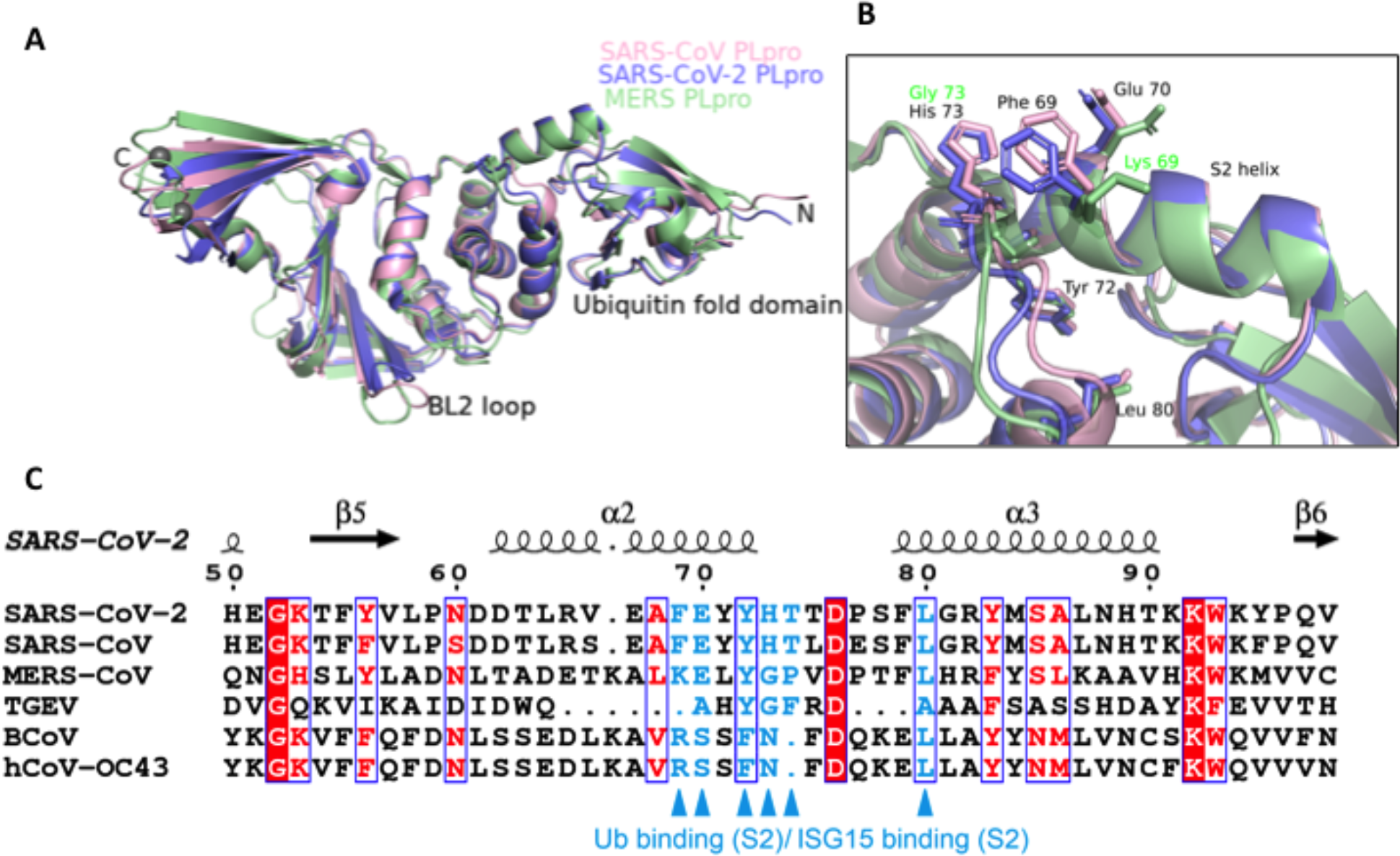
Comparison of the ISG15-S2 allosteric binding site in SARS-CoV, SARS-CoV-2 and MERS-CoV PLpro. A. Superposition of the crystal structures of SARS-CoV PLpro (PDB: 2FE8 in light pink), SARS-CoV-2 PLpro (PDB: 7NFV in slate blue) and MERS-CoV PLpro (PDB: 4RNA in light green). The zinc fingers domain, ubiquitin-fold domain and the blocking loop 2 (BL2) are the most variable regions among the three PLpro structures. B. The S2 binding ISG15 site in PLpro consisting of residues Phe 69, Glu 70, Tyr 72, His 73 and Leu 80 is conserved between SARS-CoV and SARS-CoV2. C. The sequence alignment for the region corresponding to the S2 ISG15 binding site shows that the MERS-CoV differs in critical binding residues; Phe 69 in SARS-CoV-2 PLpro is replaced by a lysine residue (F69K), Tyr 72 to a leucine residue (Y72L) and His 73 to a glycine residue (H73G).

**Fig. S4.**
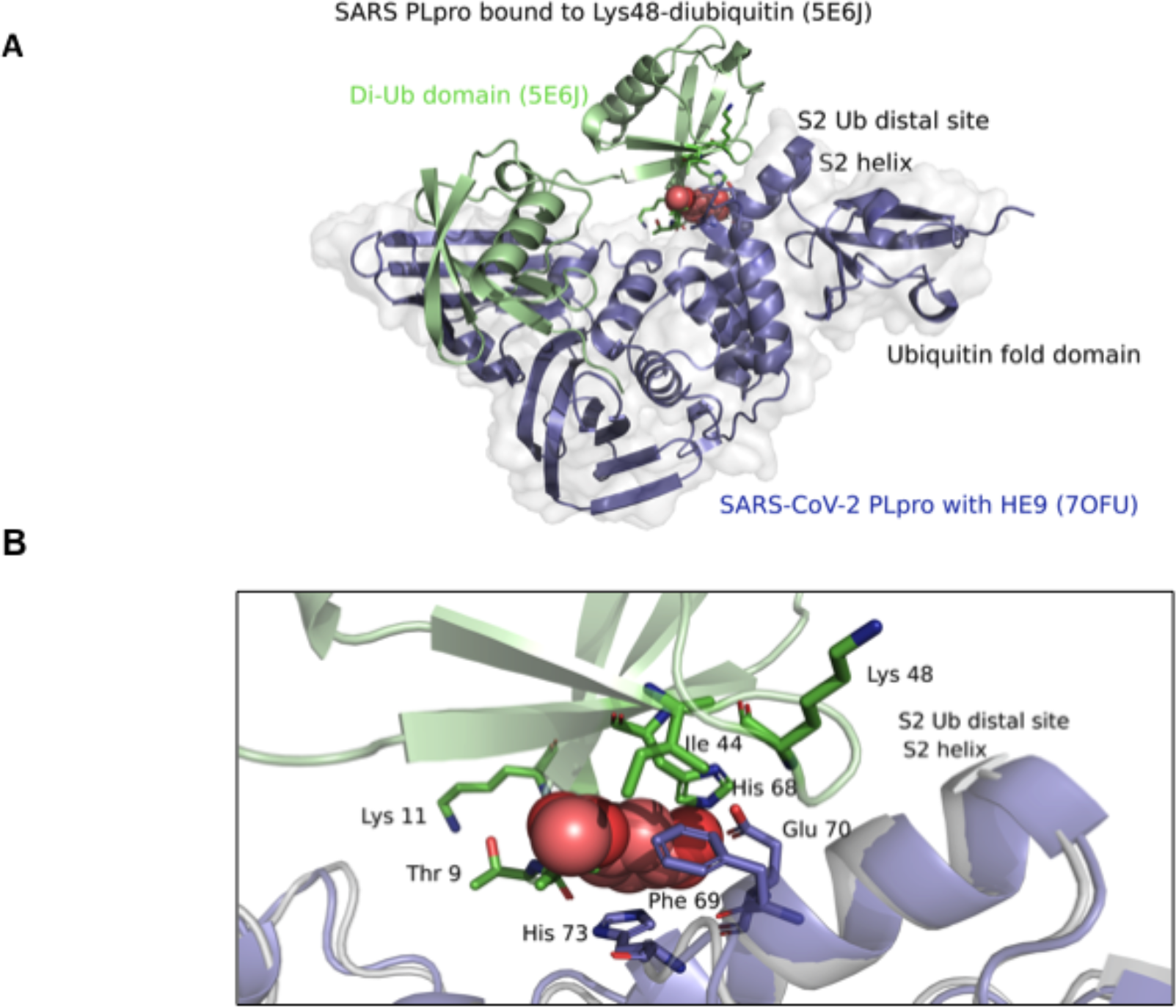
The binding of the three natural compounds disrupts interaction of the K48-linked di-ubiquitin molecule to PLpro. A. Superposition of the crystal structures of SARS-CoV PLpro complex with Lys48 linked di-Ub (PDB code 5E6J, PLpro in grey, di-Ubiquitin domain in green) with SARS-CoV-2 PLpro+HE9 (PDB code 7OFU, in slate blue). The natural compound (HE9) is represented with red spheres. B. Close-up view of the S2-Ub binding site. The di-ubiquitin molecule is shown in green cartoon representation with the interacting residues Lys 11, Ile 44 and Lys 48 shown in sticks. The bound inhibitor compound HE9 (red spheres) clearly prevents the binding of the di-ubiquitin molecule in the S2 binding site of PLpro.

**Fig. S5.**
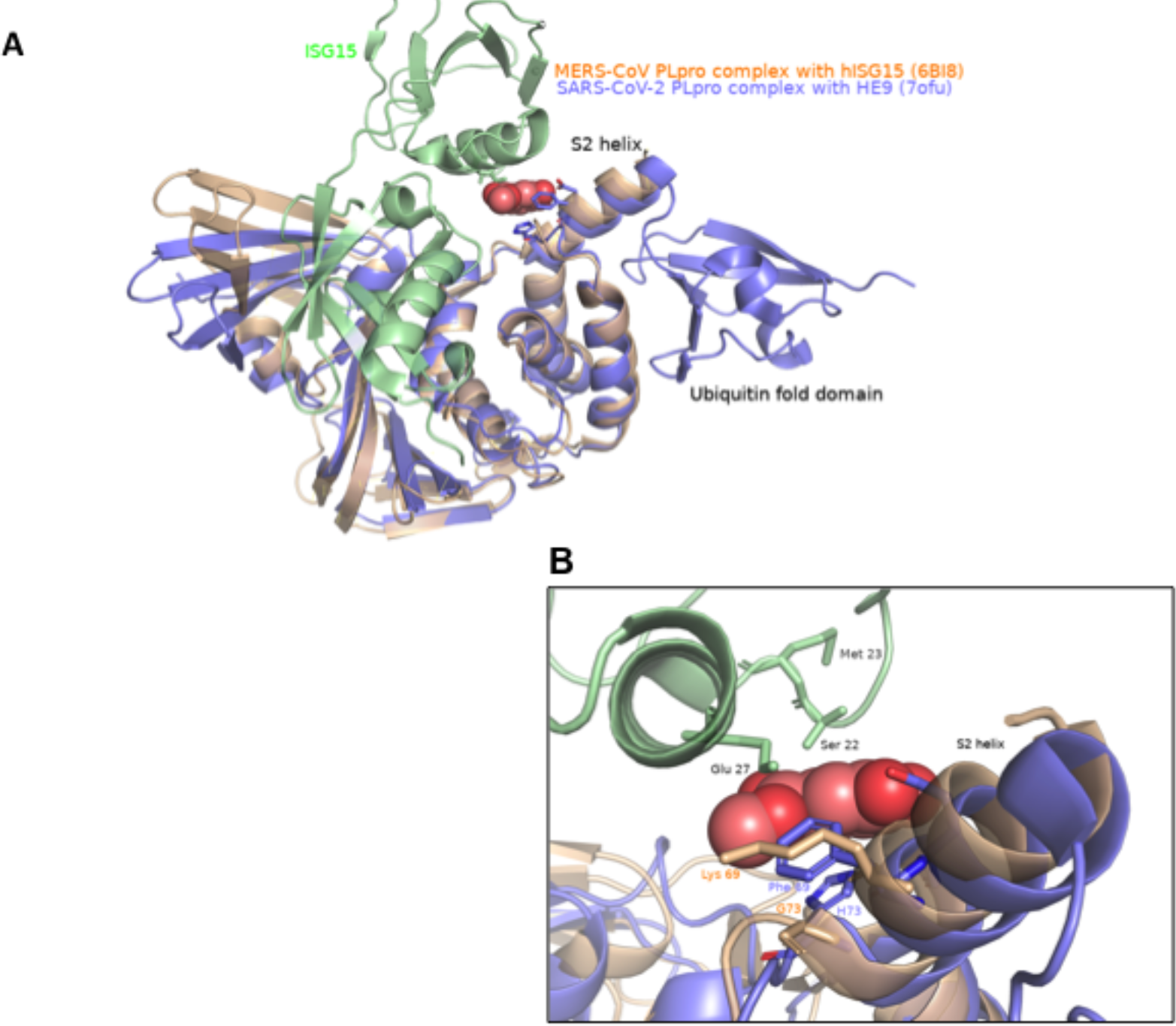
The binding of HE9 prevents interaction of the ISG15 molecule in MERS-Co-V PLpro. A. Superposition of the crystal structures of MERS-CoV PLpro complex with ISG15 (PDB code 6BI8, PLpro in wheat, ISG15 molecule in green) with SARS-Cov-2 PLpro+HE9 (PDB code 7OFU, in slate blue). The natural compound (HE9) is represented as spheres. B. Close-up view of the S2-Ub binding site. The ISG15 molecule is shown in green cartoon representation with the interacting residues Ser 22, Met 23 and Glu 27 shown in sticks. The bound inhibitor compound HE9 (red spheres) clearly blocks the binding of the ISG15 molecule to the S2 binding site of MERS-CoV PLpro. The Phe 69 to Lys and His 73 to Gly change contribute to the regional different surface charge properties of MERS-CoV PLpro.

**Fig. S6.**
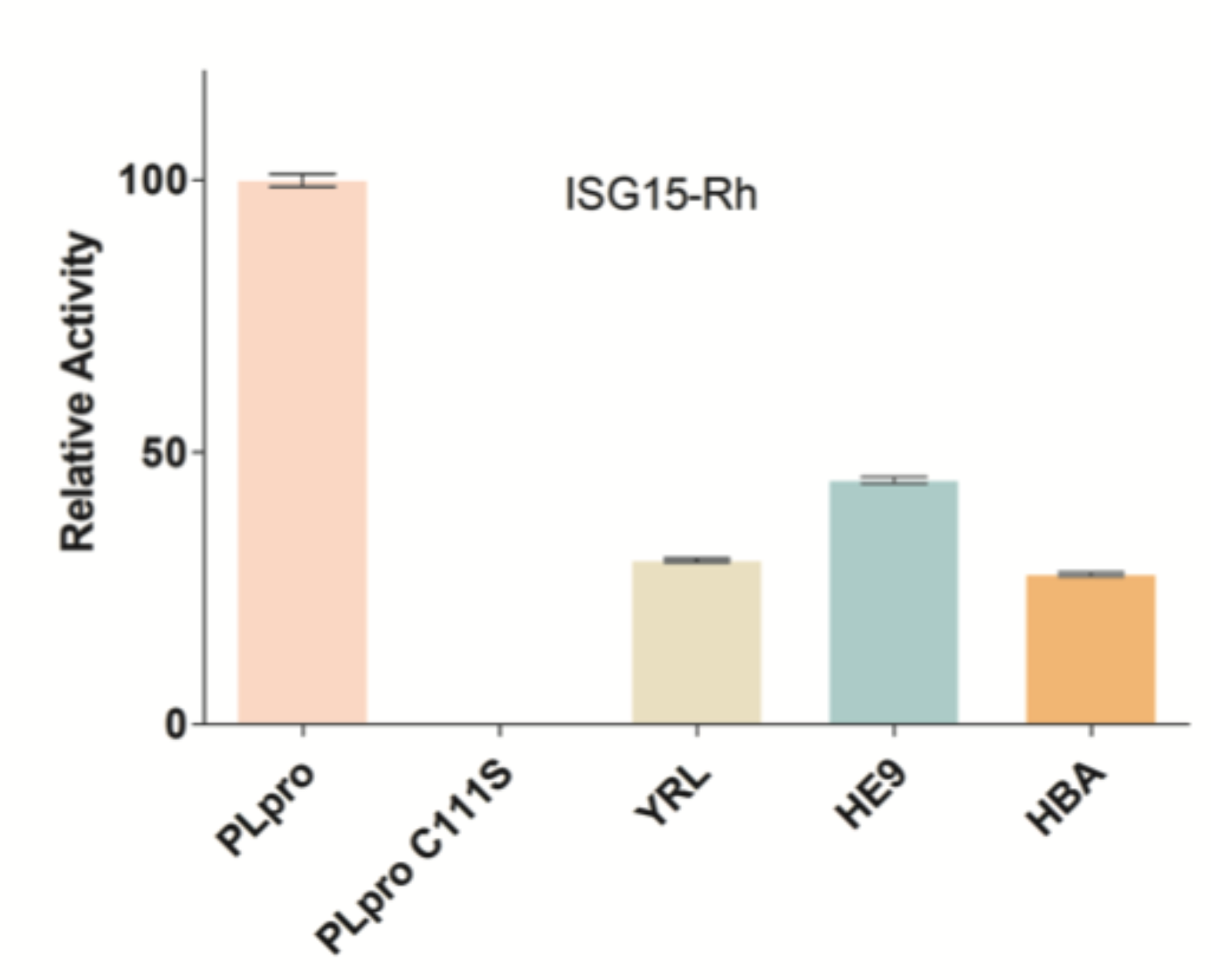
Inhibition of PLpro by the three natural compounds in deISGylation assay with ISG15-Rh substrate and comparison with catalytically inactive PLpro mutant (C111S) Wild type PLpro (WT PLpro) at 10nM concentration represents 100% of deISGylation activity and the three natural compounds YRL, HBA and HE9 at 50 µM show clear inhibition of PLpro enzymatic activity. As experimental control, catalytically inactive PLpro mutant C111S at 10nM concentration was used. ISG15-Rh at a concentration of 100nM was used as the substrate. Measured fluorescence values were blank corrected with buffer containing substrate and were measured in triplicates over 60 min with one read per minute.

**Figure S7:**
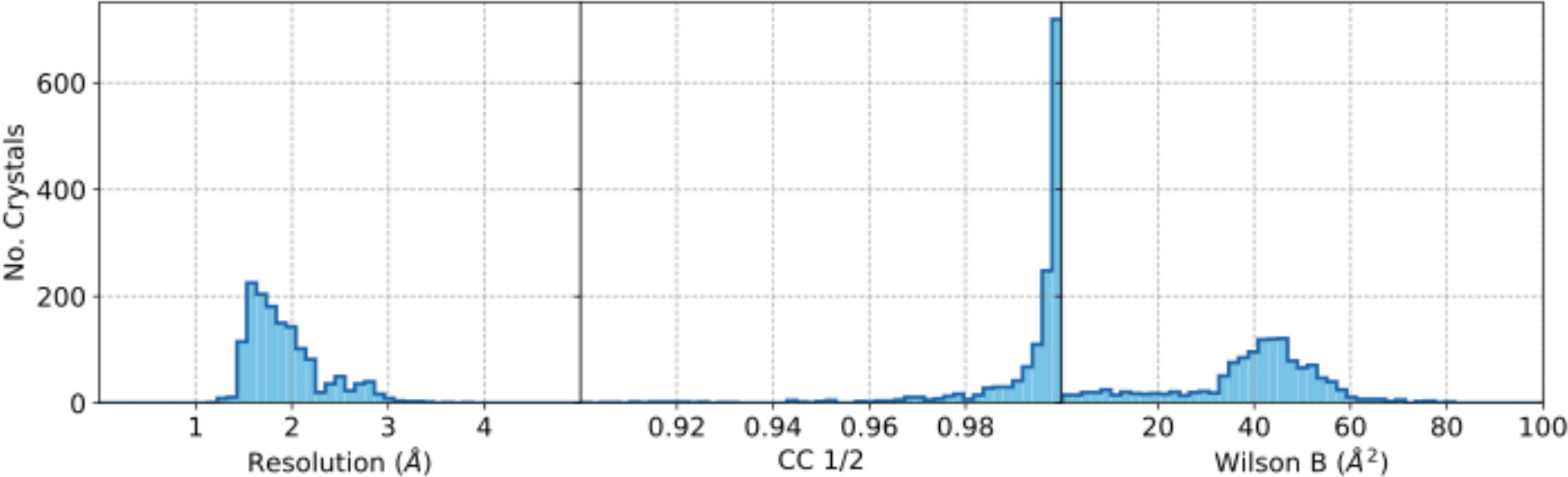
Data quality indicators of the diffraction datasets by X-ray screening of a library of 500 natural compounds. Approximately 2000 crystals of PLpro co-crystallized with the library of 500 compounds were harvested for data collection. More than one dataset per compound was collected that resulted in about 2500 datasets. A total of 1469 datasets were processed with the previously established automatic pipeline^4^. Data quality indicators, resolution (left), CC_1/2_ (middle) and Wilson B-factor (right) are shown.

**Fig. S8.**
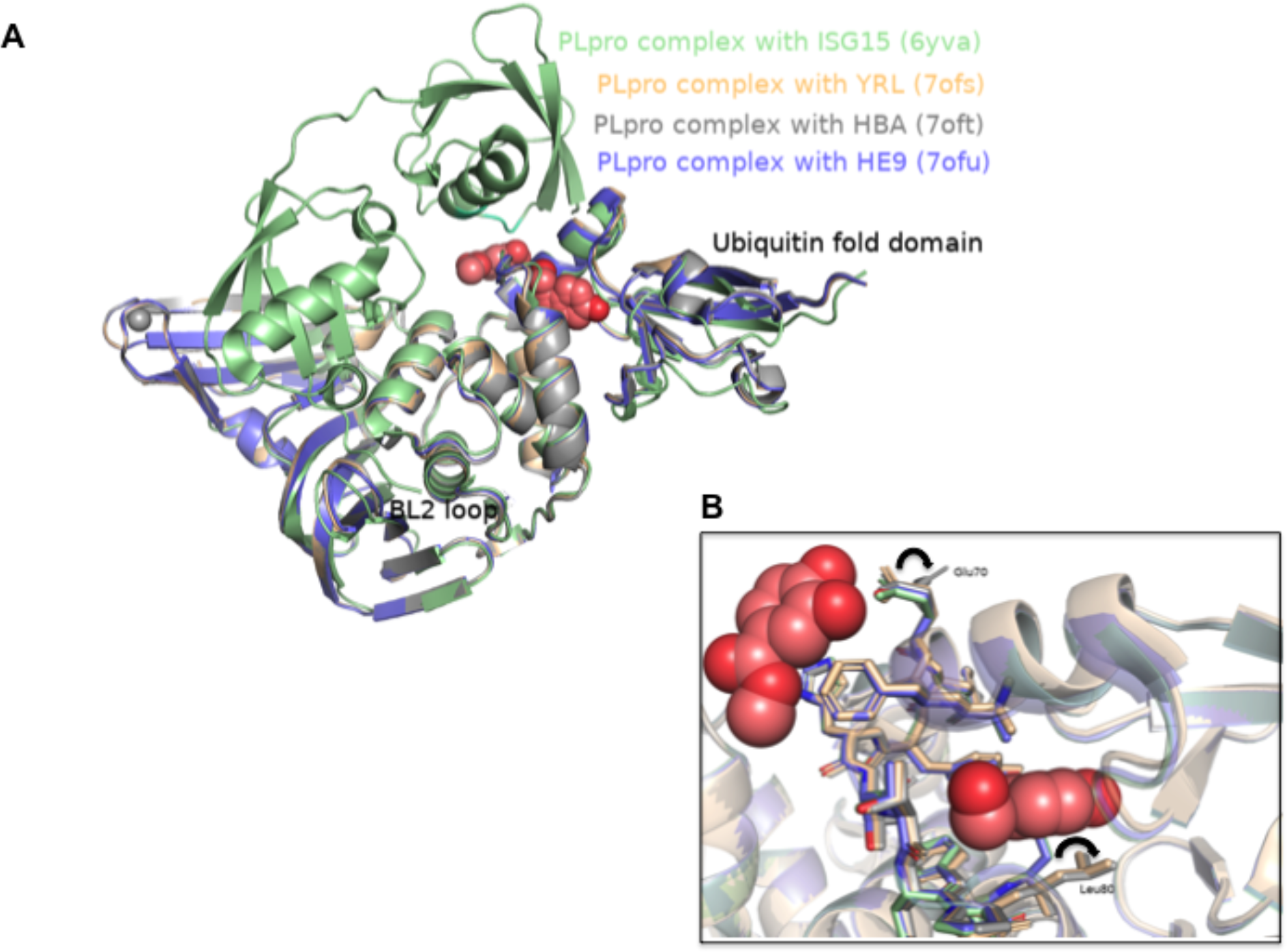
Conformational changes in PLpro by the binding of the inhibitor compounds. A. Superposition of the crystal structures of SARS-CoV-2 PLpro-C111S in complex with mouse-ISG15 (PDB code 6YVA, in light green) with SARS-CoV-2 PLpro+YRL (PDB code 7OFS, in wheat), SARS-CoV-2 PLpro+HBA (PDB code 7OFT, in grey) and SARS-CoV-2 PLpro+HE9 (PDB code 7OFU, in slate blue). It can be clearly seen that the Ubiquitin-fold domain and the BL2 loop regions are the most dynamic regions in the PLpro molecule. B. Close-up view of the ISG15 binding site. The residue Leu 80 (sticks in slate blue, apo structure) has to rotate in the complex structures to accommodate the inhibitor molecule. Glu 70 adopts a different side chain conformation when 4-hydroxybenzaldehyde (HBA) interacts with PLpro.

## Notes

### Competing Interest Statement

The authors have declared no competing interest.

